# MolDiscovery: Learning Mass Spectrometry Fragmentation of Small Molecules

**DOI:** 10.1101/2020.11.28.401943

**Authors:** Liu Cao, Mustafa Guler, Azat Tagirdzhanov, Yiyuan Lee, Alexey Gurevich, Hosein Mohimani

## Abstract

Identification of small molecules is a critical task in various areas of life science. Recent advances in mass spectrometry have enabled the collection of tandem mass spectra of small molecules from hundreds of thousands of environments. To identify which molecules are present in a sample, one can search mass spectra collected from the sample against millions of molecular structures in small molecule databases. This is a challenging task as currently it is not clear how small molecules are fragmented in mass spectrometry. The existing approaches use the domain knowledge from chemistry to predict fragmentation of molecules. However, these rule-based methods fail to explain many of the peaks in mass spectra of small molecules. Recently, spectral libraries with tens of thousands of labelled mass spectra of small molecules have emerged, paving the path for learning more accurate fragmentation models for mass spectral database search. We present molDiscovery, a mass spectral database search method that improves both efficiency and accuracy of small molecule identification by (i) utilizing an efficient algorithm to generate mass spectrometry fragmentations, and (ii) learning a probabilistic model to match small molecules with their mass spectra. We show our database search is an order of magnitude more efficient than the state-of-the-art methods, which enables searching against databases with millions of molecules. A search of over 8 million spectra from the Global Natural Product Social molecular networking infrastructure shows that our probabilistic model can correctly identify nearly six times more unique small molecules than previous methods. Moreover, by applying molDiscovery on microbial datasets with both mass spectral and genomics data we successfully discovered the novel biosynthetic gene clusters of three families of small molecules.

**Availability:** The command-line version of molDiscovery and its online web service through the GNPS infrastructure are available at https://github.com/mohimanilab/molDiscovery.

## Introduction

A crucial problem in various areas of life science is to determine which known small molecules are present/absent in a specific sample. For example, physicians are devoted to discovering small molecule biomarkers in plasma/oral/urinal/fecal/tissue samples from a patient for disease diagnosis [1] and prognosis [2]. Epidemiologists are interested in identifying small molecule disease risk factors from diet [3] and environment [4]. Ecologists are interested in characterizing the molecules produced by microbes in various microbial communities [5]. Natural product scientists need to identify all the known molecules in their sample, clearing the path towards the discovery of novel antimicrobial or antitumor molecules [6, 7].

Recent advances in high-throughput mass spectrometry have enabled collection of billions of mass spectra from hundreds of thousands of host-oriented/environmental samples [8, 9, 10, 11]. A mass spectrum is the fingerprint of a small molecule, which can be represented by a set of mass peaks (Figure 1AB). In order to identify small molecules in a sample with tens of thousands of spectra, one can either (i) de novo predict small molecule structure corresponding to mass spectra, (ii) search these mass spectra against tens of thousands of reference spectra in spectral libraries, or (iii) search these mass spectra against millions of molecular structures in small molecule databases. De novo prediction can potentially identify both known and novel small molecules. However, it is rarely used in practice due to the intrinsic complexity of small molecule structure and the low signal-to-noise ratio of mass spectral data. Spectral library search is recognized as the most reliable mass spectral annotation method. Nevertheless, current reference spectral libraries are limited to tens of thousands of molecules, and the majority of known small molecules are not represented in any reference spectral library. Furthermore, collecting mass spectra of all known small molecules individually would be expensive and time-consuming. The most frequently used strategy for small molecule identification is in-silico search of small molecule structure databases. This approach enables small molecule identification in known databases, such as PubChem [12], dictionary of natural products (DNP) [13], and AntiMarin [14]. Moreover, in-silico database search also applies to discovery of novel small molecules through genome mining [15].

**Figure 1:**
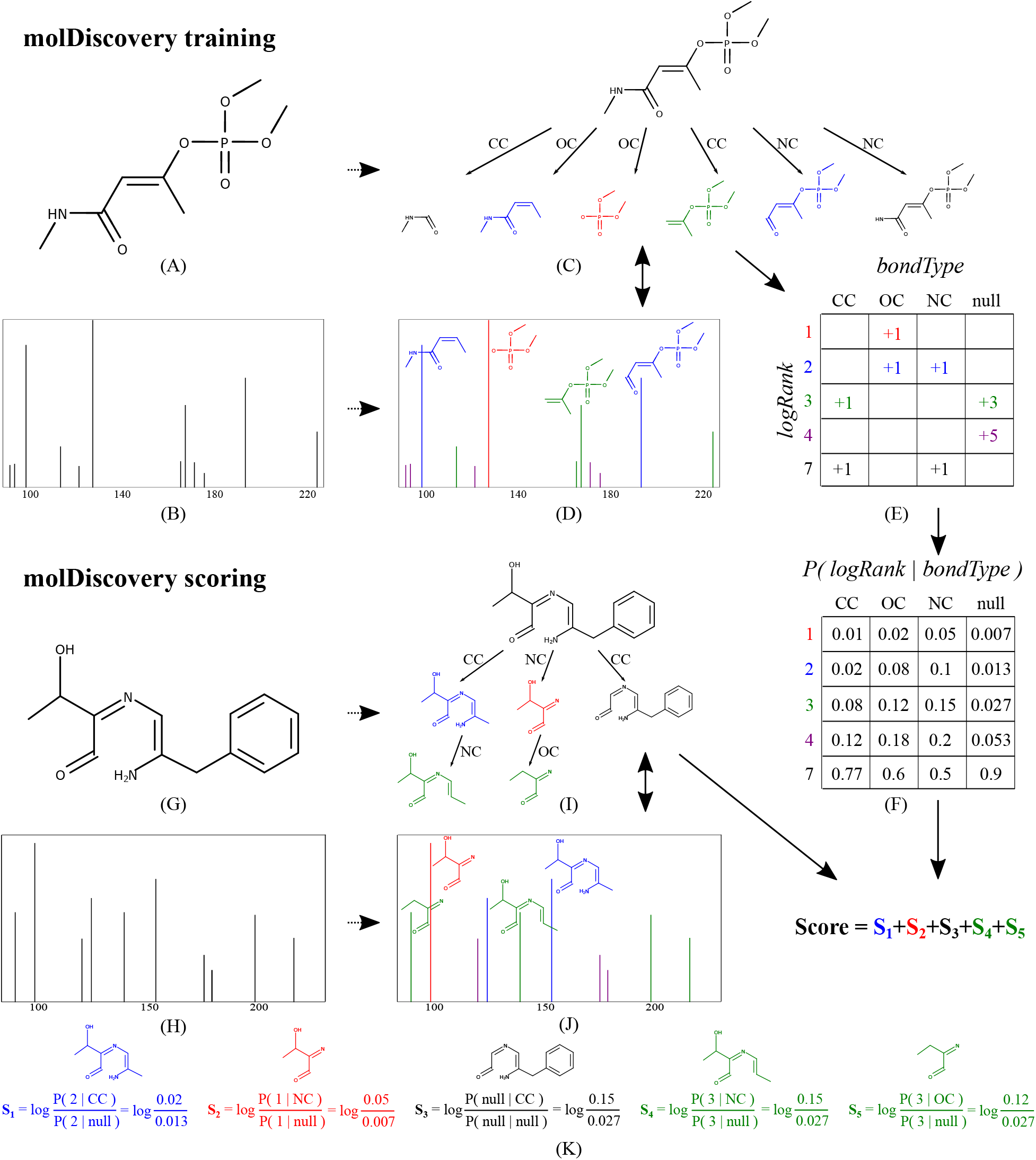
MolDiscovery Framework. Panels (A)-(F) show the training procedures of molDiscovery, while panels (G)-(K) are the scoring procedures based on the pretrained probabilistic model. (A) A reference molecule *R* in spectral library. (B) The reference spectrum of *R*. (C) The fragmentation graph of *R*. The root node represents the whole molecule, while the other nodes are fragments of it. The edges are annotated with the type of bonds where the fragmentation occur. Here, a fragment will be annotated with red, blue, green or purple if it corresponds to a mass peak in reference spectrum with *logRank* = 1, 2, 3,4 respectively (see the Methods section). (D) Annotation of the reference spectrum of *R*. A mass peak will be annotated with a fragment if its mass is identical to the mass of the fragment plus charge, within a tolerance. (E) A table counting the number of fragments observed in training data with specific *bondType* and *logRank*, referred to as count matrix. In this example, since the number of peaks is small, all the present peaks have *logRanks* 1 to 4, and the absent ones are shown with the lowest possible *logRank* = 7. Note that we allow *logRanks* between 1 – 6, corresponding to peaks with ranks between 1 – 64. The null column corresponds to the experimental peaks which cannot be explained by the fragmentation graph. (F) The probabilistic model *P*(*logRank|bondType*), which is computed by normalizing the count matrix. (G) A molecule *Q* in a chemical structure database. (H) A query spectrum. (I) The fragmentation graph of *Q*. (J) Annotation of the query spectrum with the fragmentation graph of *Q*. (K) Computation of the query spectrum match score, which is the sum of scores of all the fragments in the annotated fragmentation graph. Here we represent *P*(*logRank* = *i|bondType*) by *P*(*i|bondType*). For simplicity, *logRank_pa_*, 2-cuts columns, and rows of *logRank* = 5, 6 are not shown.

The majority of in silico database search methods are rule-based models that incorporate domain knowledge such as bond types, hydrogen rearrangement, and dissociation energy [16, 17, 18, 19, 20, 21] to predict fragmentations of small molecules and score small molecule-spectrum pairs. However, due to the complex rules involved in small molecule fragmentation, these methods are computationally inefficient for heavy small molecules. Moreover, these predictions, which are based on expert knowledge, fail to explain many of the peaks in mass spectra.

Recently, spectral libraries with tens of thousands of labelled mass spectra of small molecules (small molecule-spectrum matches) have emerged, paving the path for learning more accurate fragmentation models in order to improve sensitivity and specificity of mass spectral database search [10]. In this paper, we improve the efficiency and accuracy of small molecule identification by (i) designing an efficient algorithm to generate mass spectrometry fragmentations and (ii) developing a probabilistic model to identify small molecules from their mass spectra. Our results show that molDiscovery greatly increases the accuracy of small molecule identification, while making the search an order of magnitude more efficient. After searching 8 million tandem mass spectra from the Global Natural Product Social molecular networking infrastructure (GNPS) [10], molDiscovery identified 56,971 small molecules at 0% false discovery rate (FDR), an eight times increase compared to existing methods. On a subset of the GNPS repository with known genomes, molDiscovery correctly linked 20 known and three novel biosynthetic gene clusters to their molecular products.

## Results

### Outline of molDiscovery pipeline

The molDiscovery pipeline starts by (i) constructing metabolite graphs and (ii) generating fragmentation graphs. For the latter, molDiscovery uses a new efficient algorithm for finding bridges and 2-cuts in the metabolite graphs. Afterwards, molDiscovery proceeds with (iii) learning a probabilistic model for matching fragmentation graphs and mass spectra (Figure 1A-E), (iv) scoring small molecule-spectrum pairs (Figure 1F-K), and (v) computing FDR.

In the past we introduced Dereplicator+ [22], a database search method for identification of small molecules from their mass spectra, that follows a similar series of steps to search mass spectra against chemical structures. However, Dereplicator+ uses a brute-force method for fragmentation graph construction, and the naive shared-peak-count for scoring.

### Datasets

The GNPS spectral library contains 8,964 small molecule-spectrum matches [10]. After removing spectra corresponding to duplicate molecules, we obtain 4,781 non-redundant spectra, which are used as training and testing data for molDiscovery. The MassBank of North America (MoNA) contains 86,209 positive mode LC-MS/MS spectra. After filtering for unique spectra using SPLASH [23] and isolating singly-charged molecule-spectrum matches, spectra from 3,278 unique molecules are extracted (determined using RDKit [24] canonicalized SMILES). These two spectral libraries are searched against the combined natural product structure database (CombinedDB) of DNP (77,057 non-redundant molecules) [13] and AntiMarin (60,908 molecules) [14].

Furthermore, we benchmark molDiscovery against Dereplicator+ [22] using 46 high-resolution GNPS spectral datasets (Supplementary Table S1) collected from various environments (about 8 million mass spectra in total). These spectra are searched against a combined structure database (AllDB) of 719,958 compounds from DNP, AntiMarin, UNPD [25], HMDB [26], LMSD [27], FooDB [28], NPAtlas [29], KEGG [30], DrugBank [31], StreptomeDB [32], GNPS spectral library, MIBiG [33], PhenolDB [34] and Quorumpeps [35] databases.

### Preferential fragmentation patterns in mass spectrometry

MolDiscovery learned a probabilistic model (see Methods) that reveals the preferences in mass spectrometry fragmentation. First, mass spectrometry fragmentation has preference for bond type *bondType*. Figure 2A shows that the breakage of *bondTypes* NC and OC lead to fragments with higher rank (i.e. *logRanks* closer to 1) than CC bond, indicating that they are more likely to be broken by mass spectrometry. In addition, bridges (NC, OC and CC) tend to generate fragments with higher *logRank* than 2-cuts (for example, (NC_NC, NC_OC, CC_OC), suggesting bridges are more likely to be fragmented than 2-cuts. Note that the *logRank* distribution of fragments due to the 2-cut CC_CC is similar to the null model distribution (random noise), which implies that in practice the fragmentation of CC_CC 2-cut rarely happens.

**Figure 2:**
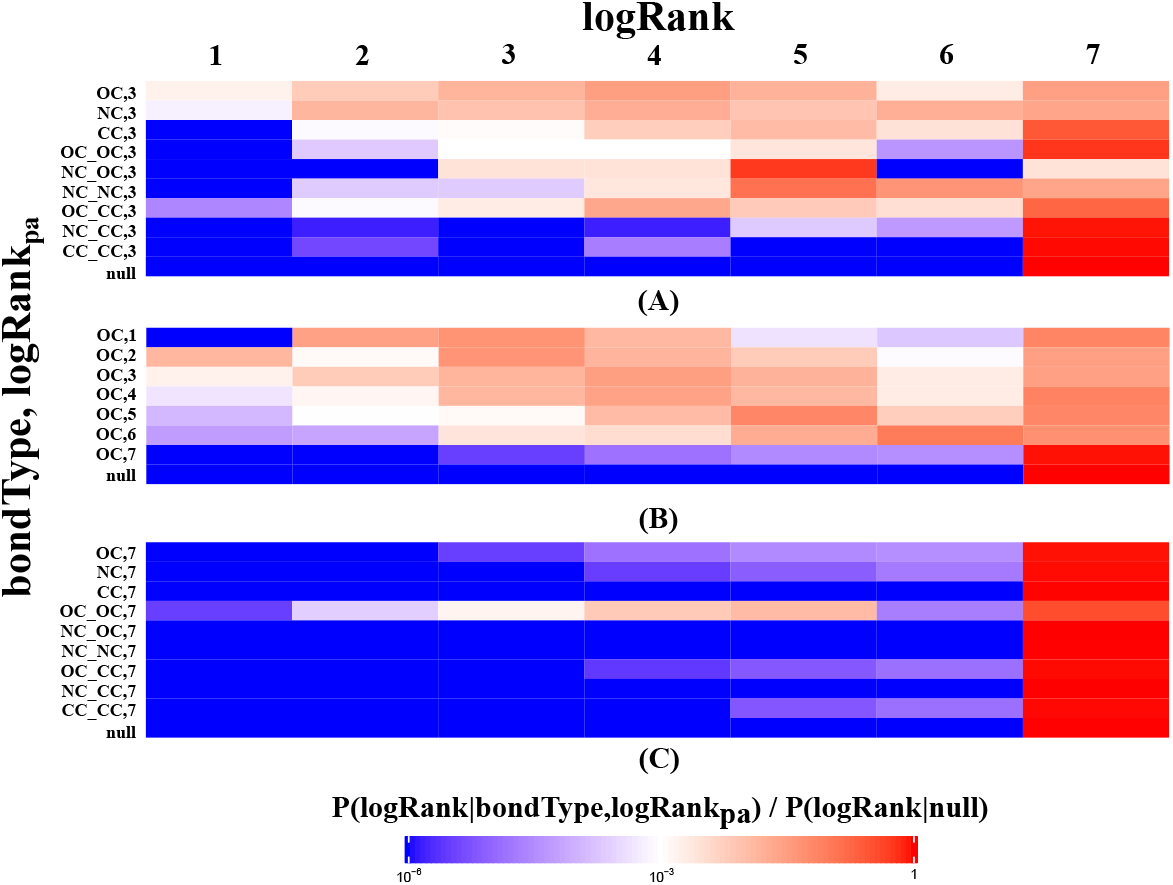
Heatmap of *P*(*logRank|bondType, logRank_pa_*). Each row represents bond type *bondType* and parent’s intensity rank *logRank_pa_* of a fragment. The row ‘‘null” refers to the null distribution *P*(*logRank|null*). Each column represents the *logRank* of a child fragment. When *logRank_pa_* is 0, it means the parent is the root (precursor molecule). (A) *logRank* distribution over different *bondTypes* for *logRank_pa_* = 3. (B) *logRank* distribution of OC bond over different *logRank_pa_*. Since there is only one fragment in the fragmentation graph that could have *logRank* = 1, the parent fragment of such fragment can not generate another fragment with *logRank* = 1, hence *P*(1|*NC*, 1) = 0. (C) *logRank* distribution over different *bondTypes* for *logRank_pa_* = 7. Supplementary Figure S1-2 shows the complete heatmaps for charge +1 and +2 respectively.

In addition, parent fragments with high ranks are more likely to produce high-rank children fragments, while fragments with low ranks are similar to random noise. For example, for fragments produced by the breakage of OC bond (Figure 2B), if their parents have high ranks (1, 2 or 3), they are likely to be higher rank than those with lower-rank parents (6, 7). Moreover, molDiscovery automatically learns to discard fragments with low rank parents which are not informative for predicting mass spectrometry fragmentations. The distribution of *logRank* for these fragments is similar to noise (null distribution, Figure 2C). Therefore, the log-likelihood scores for such frag-ments are close to zero.

### Benchmarking the efficiency of fragmentation graph construction algorithms

We compared the performance of the fragmentation graph construction algorithms in molDiscovery and Dereplicator+. For both algorithms, we allow for at most two bridges and one 2-cut. The algorithms are benchmarked on CombinedDB. Figure 3A and Supplementary Figure S3A show the average fragmentation graph construction time for small molecules in different mass ranges. The results show that the average fragmentation graph construction time for Dereplicator+ grows exponentially as the mass of small molecules increase, while the construction time only grows linearly for molDiscovery. When the mass is greater than 600 Da, molDiscovery is two orders of magnitude faster than Dereplicator+.

**Figure 3:**
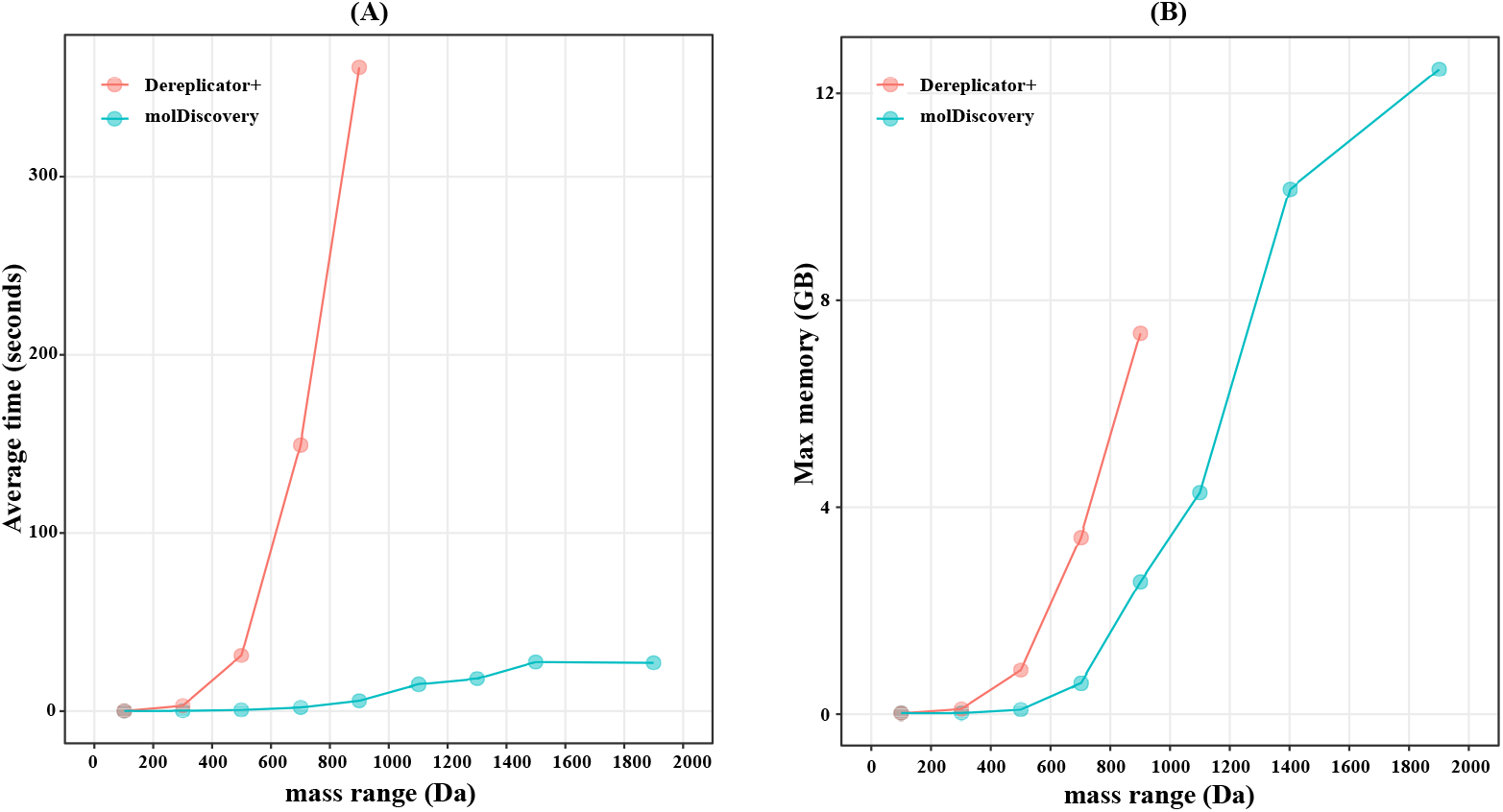
Benchmarking the efficiency of fragmentation graph construction algorithms. We compare (A) the average fragmentation graph construction time and (B) maximum memory consumption of molDiscovery with Dereplicator+ on the AntiMarin database over molecules of various masses. Note that we randomly selected 200 molecules from the AntiMarin database for each 200 Da mass window to compute average running time per molecule as well as maximum memory usage. For cases exceeding the memory limitation of 15 GB, no data point is shown.

Figure 3B and Supplementary Figure 3B compare the maximum memory consumption of molDiscovery with Dereplicator+. For both chemical structure databases, Dereplicator+ only works for molecules with masses less than 1000 Da, while molDiscovery can handle molecules with masses up to 2000 Da.

### Benchmarking database search accuracy of molDiscovery

The accuracy of database search by molDiscovery is compared to four state-of-the-art methods on the non-redundant GNPS spectral library. First we split the spectra into five batches, each with 956 and one with 957 tandem mass spectra. In order to evaluate their performance on larger chemical structure databases, we combined the molecules in the non-redundant GNPS spectral library with non-redundant DNP (77,057 molecules). Then we performed 5-fold cross validation for molDiscovery against this combined database of chemical structures. The database search results show that molDiscovery on average can correctly identify 74.2% small molecules in the test dataset as top ranked identifications in comparison to 63.6% for Dereplicator+, and lower rates for MAGMa+ [37], and CFM-ID [38] (Figure 4A). CSI:FingerID [39] achieves an average top ranked identification rate of 73.5%. However, of the 4,781 molecules tested 3,042 molecules appear in the CSI:FingerID positive mode training dataset. If we only evaluate the 1,739 compounds that are not present in the CSI:FingerID training dataset, molDiscovery achieves 73.9% and 92.2% accuracy for top 1 and top 10 identifications respectively, while CSI:FingerID achieves 66.1% and 71.0% accuracy (Figure 4B). Supplementary Figure 4 shows the top-k accuracies of the methods in the five cross validation batches.

**Figure 4:**
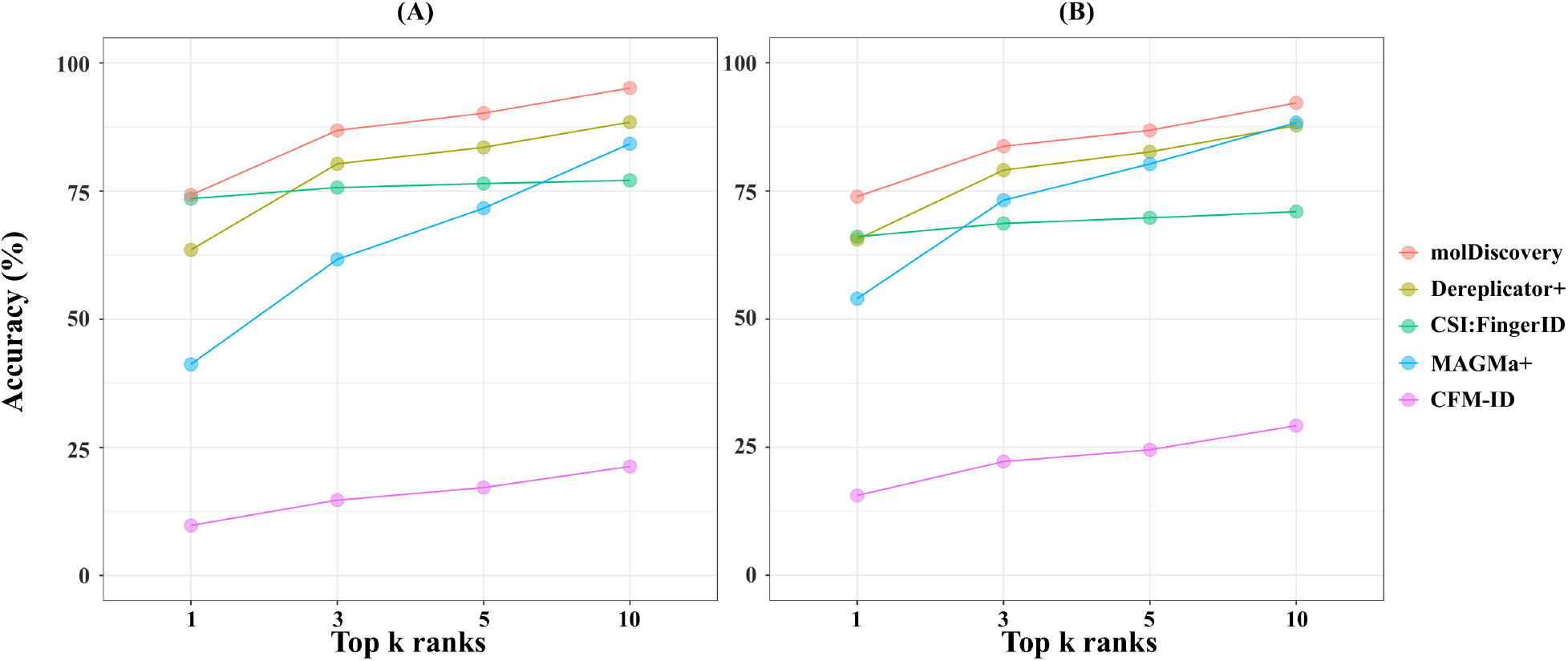
Benchmarking the database search accuracy on (A) GNPS spectral library (4,781 molecules) and (B) GNPS spectral library excluding CSI:FingerID training data (1,739 molecules). For each spectrum, we rank the matching scores given by molDiscovery, Dereplicator+, MAGMa+ 1.0.1, CFM-ID 2.0, and CSI:FingerID 1.4.3, (from SIRIUS 4.4.2 [36]). If there is a tie, we assign the average rank to all the matches in the tie. Then we check if the correct molecule-spectrum match is among the top *k* (*k* = 1, 3, 5,10) matches. The x-axis is the rank *k*, and the y-axis is the mean proportion of correct molecule-spectrum matches in the test datasets that are among the top *k* matches.

MolDiscovery was also the most efficient method and used only 217 MB of memory as its peak (Table 1) making it seven times faster and two orders of magnitude more memory efficient than Dereplicator+. MolDiscovery was three orders of magnitude faster than CSI:FingerID, CFM-ID, and MAGMa+.

**Table 1:** Runtime and memory usage of various *in silico* search tools. Reported performance is for a search of 4,781 spectra against DNP+GNPS (81,838 compounds). If supported by the software the chemical database was pre-processed before the search was run.

| Software | Runtime (d:hh:mm) | Memory Usage (GB) |
| --- | --- | --- |
| molDiscovery | 0:00:28 | 0.22 |
| Dereplicator+ | 0:03:23 | 25.46 |
| CFM-ID | 8:00:44 | 0.041 |
| CSI:FingerID | 7:07:26 | 3.13 |
| MAGMa+ | 3:14:27 | 1.87 |

### Evaluating molDiscovery sensitivity to platform variation

In order to determine if the probabilistic scoring method described is robust to changes in mass spectrometry platform, a subset of annotated spectra from MoNA dataset including 1,639 HCD-fragmented and 1,639 CID-fragmented spectra were used for evaluation. Two models were trained using the GNPS spectra augmented with either the HCD or CID fragmented MoNA dataset, and then tested on both fragmentation modes. Searches were run against a combined chemical structure database consisting of 3,278 MoNA, 77,057 non-redundant DNP, and 4,781 GNPS molecules. It was shown that the platform on which training data was collected did not affect performance of molDiscovery (Figure 5).

**Figure 5:**
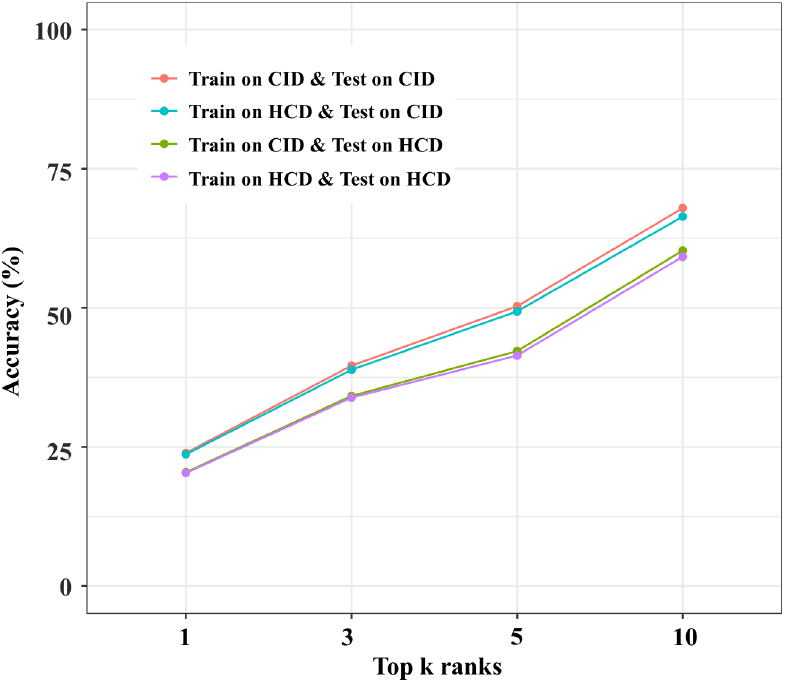
Accuracy of molDiscovery on tandem mass spectra of different fragmentation modes. MolDiscovery models were trained on the GNPS spectra augmented with MoNA spectra labeled with either CID or HCD fragmentation modes. The two models are then tested on both CID and HCD data. Matches were ranked according to score. The fraction of spectra for which the correct molecule was in the top *k* (*k* = 1, 3, 5,10) matches are shown along the *y*-axis.

### Performance on doubly-charged spectra

Only 163 spectra from MoNA dataset are annotated as doubly-charged. In light of the small amount of data we reserved the MoNA dataset as our test dataset, and used high confidence doubly-charged Dereplicator+ identifications of the GNPS datasets as the training data (813 spectra and 180 unique molecules at cuttoff score of 15). Since this training data is still small, we bootstrapped doubly-charged parameters using the singly-charged model we previously trained. The training process updates the model parameters according to the doubly-charged training data. Despite the small training dataset, molDiscovery outperformed Dereplicator+ on search of 163 doubly-charged spectra against non-redundant DNP and GNPS chemical database augmented with MoNA molecules (Figure 6).

**Figure 6:**
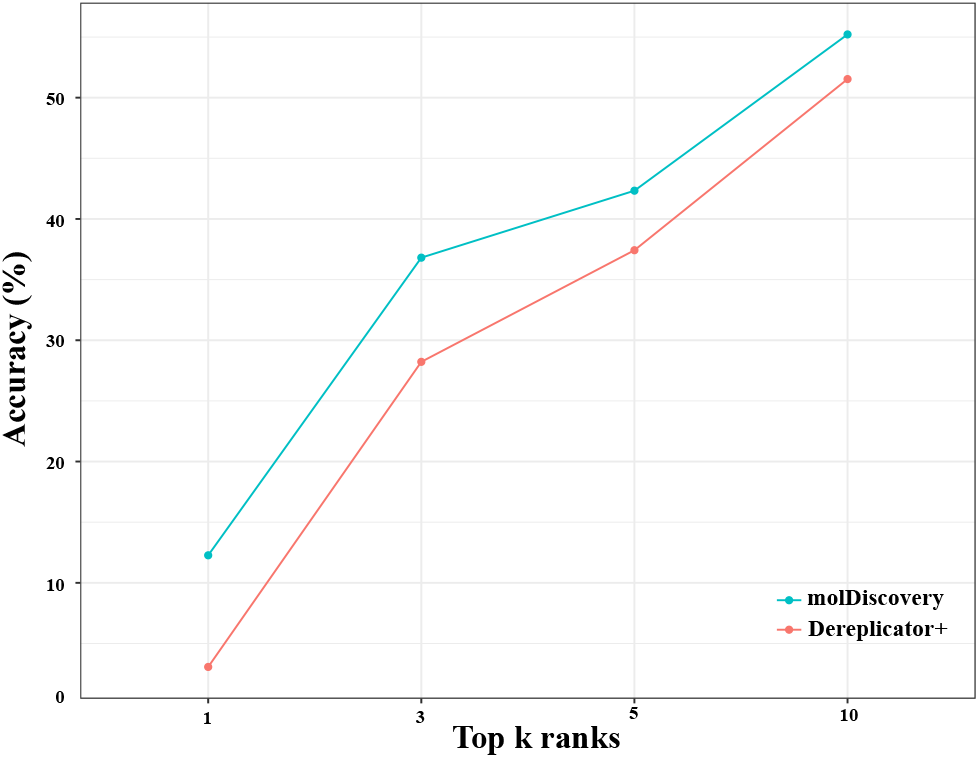
Accuracy of Dereplicator+ and molDiscovery on 163 doubly-charged spectra from MoNA. The molDiscovery model was initialially trained on 4,781 singly-charged spectra from GNPS and further trained on 813 spectra identified as doubly-charged spectra by Dereplicator+.

### MolDiscovery performance on different compound classes and mass ranges

To evaluate how well molDiscovery performs on different compound classes, we used ClassyFire [40] to annotate the non-redundant GNPS spectral library. Figure 7A shows that molDiscovery performs as well as or better than all the other tools on 15 out of 17 superclasses. The two superclasses for which molDiscovery has lower accuracy than other methods, *Homogeneous non-metal compounds* and *Hydrocarbon derivatives*, had only five and two annotated compounds respectively (Supplementary Figure S5A). Similarly, molDiscovery performs equally or better than the competing methods on compounds from nearly all mass ranges (Figure 7B). It is worth noting that the accuracy of CSI:FingerID drops a lot on the compounds that are not in its training data (Supplementary Figure S6).

**Figure 7:**
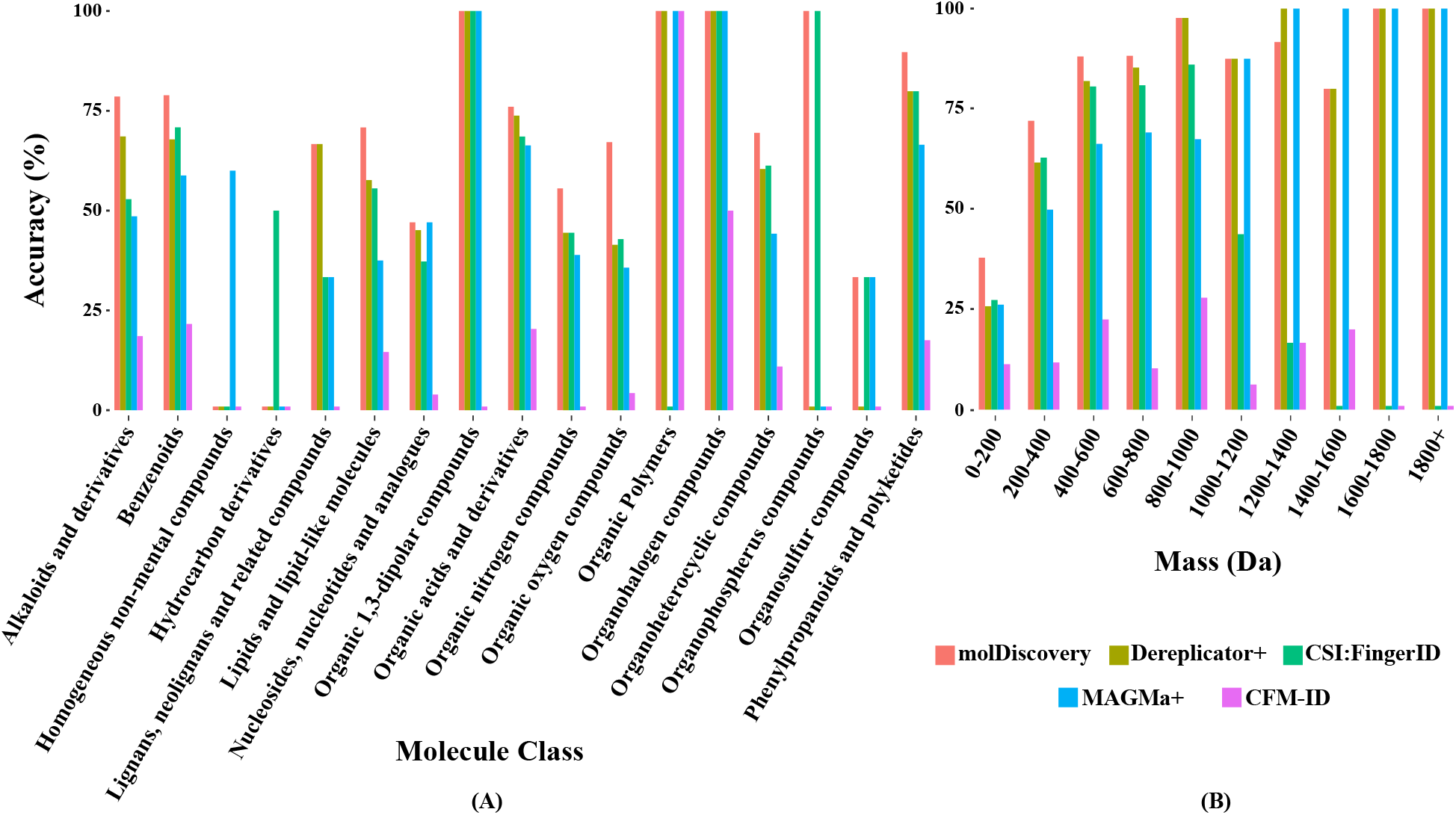
Database search accuracy of molDiscovery, Dereplicator+, CSI:FingerID, MAGMa+, and CFM-ID on molecules of (A) different superclasses from ClassyFire [40] and (B) different mass ranges on GNPS spectral library compounds excluding CSI:FingerID training data.

### Performance on large-scale spectral datasets

We benchmarked the performance of molDiscovery against Dereplicator+ on 45 GNPS datasets containing a total of 8 million tandem mass spectra (Supplementary Table S1). In contrast to the GNPS spectral library where all the spectra are annotated and their molecules are known, the spectra in the GNPS spectral datasets are not annotated, and they could correspond to novel or known molecules. Figure 8 shows the numbers of small molecule-spectrum matches and unique compounds from the AllDB database identified by molDiscovery and Dereplicator+ at various FDR levels. At the strict 0% FDR level, molDiscovery identified eight times more spectra (56971 versus 7684) and six times more unique compounds (3185 versus 542) than Dereplicator+. MolDiscovery search took 34 days on 10 threads, and CSI-FingerID, CFM-ID and MAGMa+ are not benchmarked, as the searches would have taken hundreds of years. The top 100 identifications by molDiscovery is shown in Supplementary Table S2.

**Figure 8:**
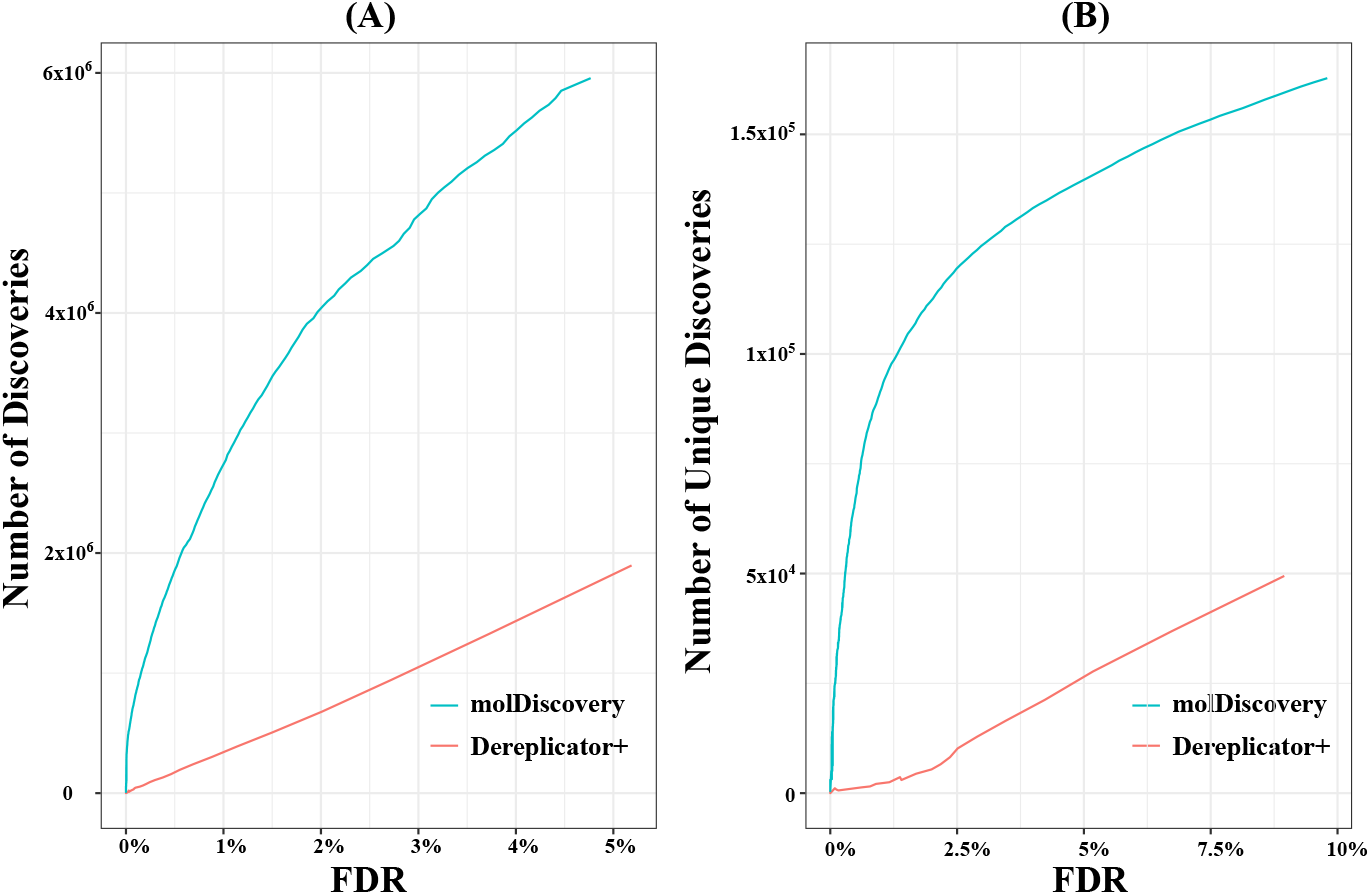
Performance of molDiscovery and Dereplicator+ on GNPS spectral datasets. The curves show the number of (A) small molecule-spectrum matches and (B) unique compounds identified by molDiscovery and Dereplicator+ in the search of 8 million spectra from 46 GNPS spectral datasets against 719,958 compounds of AllDB at different FDR levels.

### Validating molDiscovery identifications using a literature search

We benchmarked molDiscovery against Dereplicator+ on top 100 identifications from GNPS datasets by performing literature search. For this task, we used the extensively studied GNPS dataset MSV000079450 (~ 400, 000 spectra from *Pseudomonas* isolates) [41, 42]. Out of the top 100 small molecule-spectra matches reported by molDiscovery, 78 correspond to compounds having *Pseudomonas* origin based on taxonomies reported for molecules in AntiMarin database. The second largest genus among the identifications (20 out of 100) is *Bacillus*. The molecule-spectrum matches from *Bacillus* are likely to be true positives as the dataset is known to be contaminated with *Bacillus* species [42]. While the top 100 identifications from Dereplicator+ also contains 20 *Bacillus* matches, the number of hits related to *Pseudomonas* species is 62, 25% lower than molDiscovery. 18 identifications of Dereplicator+ are annotated as having fungi origin, which are likely false positives.

### MolDiscovery links small molecules to their biosynthetic gene clusters (BGCs)

We cross-validated molDiscovery and genome mining by searching 46 microbial datasets (922 unique microbial strains, 8,013,433 spectra) that contained both tandem mass spectrometry data and genomic data. MolDiscovery successfully identified 20 molecules in microbial isolates that contained their known BGCs (Table 2.)

**Table 2:**
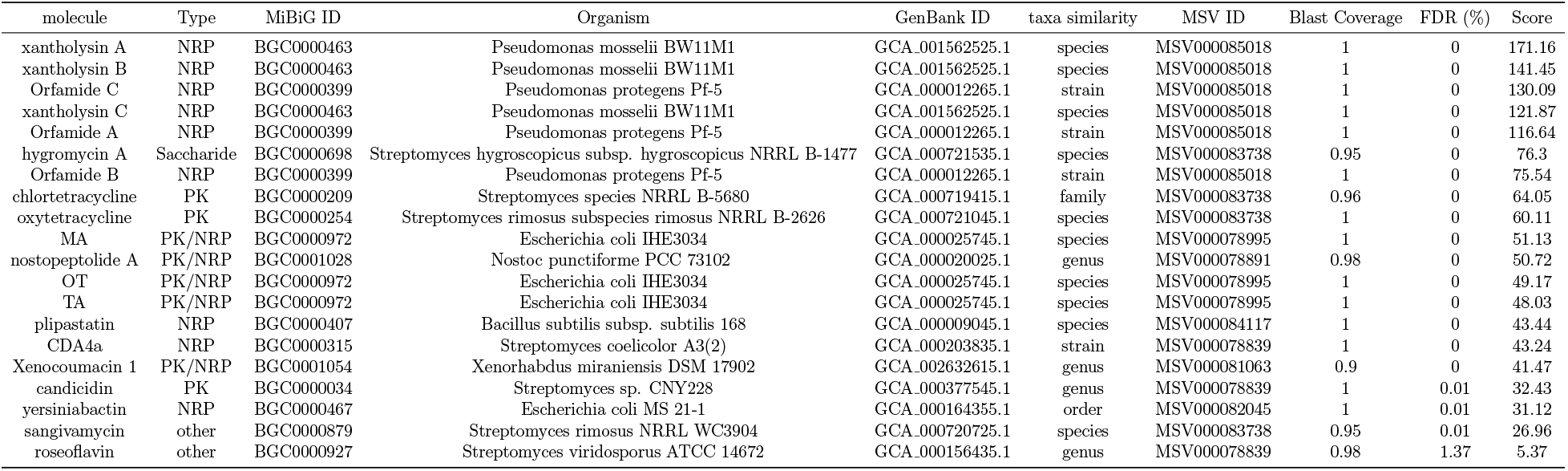
MolDisovery identified 20 molecules in bacterial isolates that contained their known BGCs. A genome is reported to contain a known BGC from the MIBiG database [33] if the genome has total BLAST hit [43] to the BGC with at least 90% coverage and 70% identity. Taxa similarity means similarity between the taxon of source organism and the organism reported in MiBIG. PK stands for polyketides, MA stands for N-myristoyl-D-asparagine, OT stands for (R)-N1-((S)-5-oxohexan-2-yl)-2-tetradecanamidosuccinamide, and TA stands for cis-7-tetradecenoyl-D-asparagine.

Moreover, molDiscovery successfully discovered novel BGCs for three small molecule families from streptomyces dataset MSV000083738 [44, 45]. MolDiscovery search results for this dataset are available at https://github.com/mohimanilab/molDiscovery. MolDiscovery identified dinghu-peptins A-D [46] in *Streptomyces* sp. NRRL B-5680 and several other *Streptomyces* strains (Supplementary Table S2). AntiSMASH [47] revealed a non-ribosomal peptide (NRP) biosynthetic gene cluster in the genome of this strain, with seven adenylation domains that are highly specific to the amino acid residues of dinghupeptin molecules [48] (Supplementary Figure S7).

MolDiscovery detected lipopeptin family and neopeptin family molecules in *Streptomyces hygroscopicus* NRRL B-1477 and *Streptomyces rimosus* NRRL WC3874. Genome mining revealed an NRP BGC in *S. hygroscopicus* NRRL B-1477 with seven adenylation domains, which are highly specific to the residues of neopeptin and lipopeptin (Supplementary Figure S8). It has been reported that these two families are structurally similar [49]. Thus, this BGC could be responsible for the production of both families.

In addition, molDiscovery identified longicatenamycin family of cyclic hexapeptides in *Streptomyces rimosus* NRRL B-8076 and several other *Streptomyces* strains. Genome mining of the NRRL B-8076 strain revealed a BGC highly specific to longicatenamycin (Figure 9). MolDiscovery also identified cyclic hexapeptide families nicrophorusamide (A and B) and noursamycin (A, D and E) in related *Streptomyces* strains (Supplementary Table S2), suggesting the same BGC might be responsible for their production. Dereplicator+ and CSI-FingerID fail to discover longicatenamycin at 1% FDR threshold.

**Figure 9:**
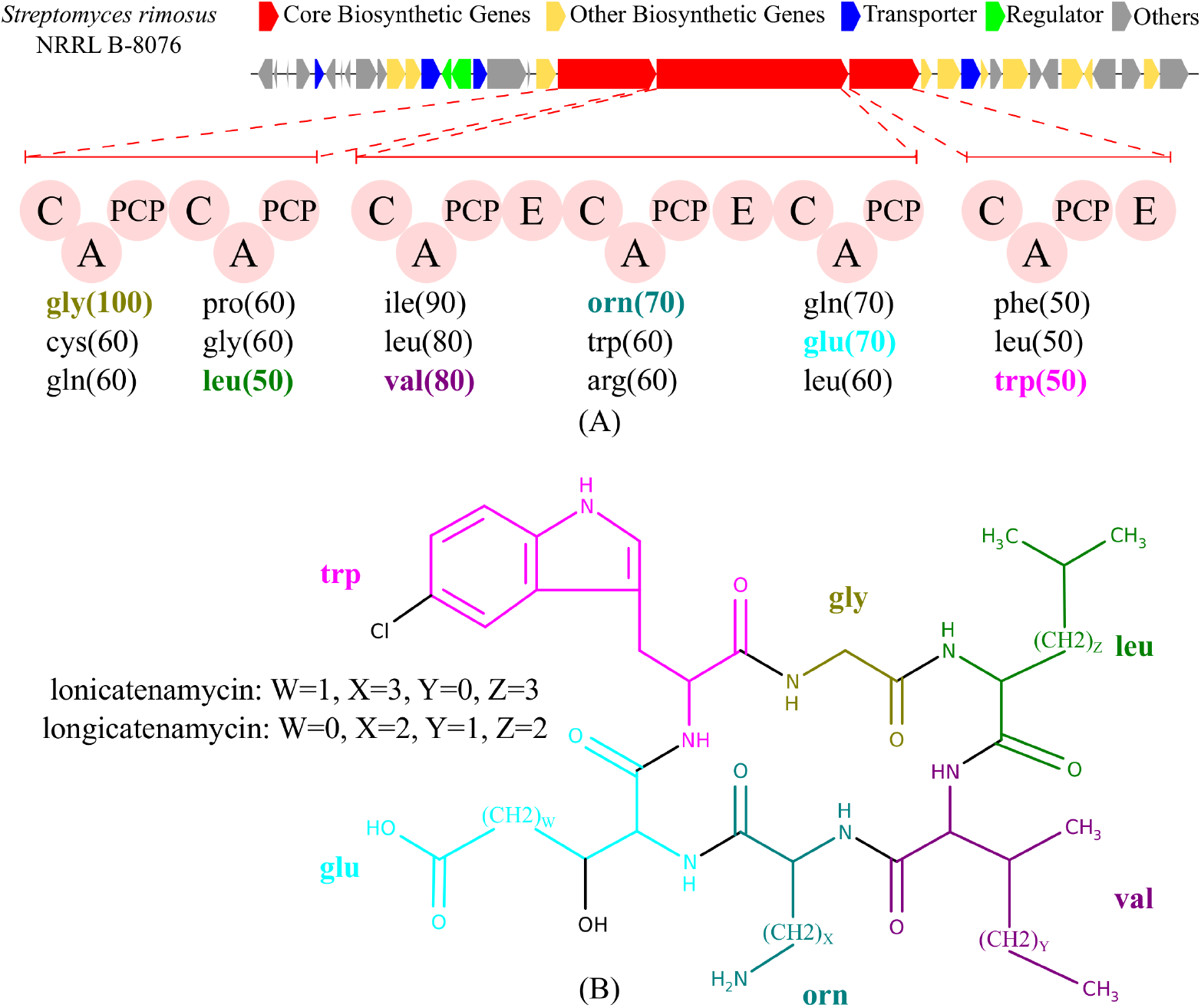
(A) Biosynthetic gene cluster of longicatenamycin family. MolDiscovery identified longicatenamycin variants at FDR 0% in *Streptomyces rimosus* NRRL B-8076. After searching its genome using antiSMASH, we detected an NRP BGC with adenylation domains which show high specificity to the amino acids residues of longicatenamycin. (B) Molecular structures of the two longicatenamycin variants identified by molDiscovery at FDR 0%.

## Discussion

With the advent of high-throughput mass spectrometry, large-scale tandem mass spectral datasets from various environmental/clinical sources have become available. One of the approaches for annotating these datasets is in silico database search. Currently the existing methods for in silico database search of tandem mass spectra are rule-based. These methods can not scale to searching molecules heavier than 1000 Da and fail to explain many of the fragment peaks in tandem mass spectra.

MolDiscovery introduces an efficient algorithm to construct fragmentation graphs of small molecules, enabling in silico database search for molecules up to 2000 Da. MolDiscovery is more efficient than the state-of-the-art methods in terms of run-time and memory consumption. Furthermore, by training on the reference spectra from the GNPS spectral library, molDiscovery learned a probabilistic model that significantly improves the accuracy of database search. MolDiscovery is shown to be robust against dissociation technique, and it performs well on singly and doubly charged spectra.

Small molecule fragmentation is a complex process that depends not only on the type of fragmented bonds, but also on local/global features of small molecules such as moiety. Currently, molDiscovery ignores all these features. Recent advances in graph-based machine learning has enabled representing complex small molecule structures with continuous vectors, making it feasible to incorporate local/global structural information into the prediction of fragmentation, potentially leading to more accurate fragmentation models. MolDiscovery paves the way for these more sophisticated approaches by collecting larger training data of small molecule-spectrum matches through the search of millions of spectra.

## Methods

### Constructing fragmentation graphs of small molecules

MolDiscovery first constructs a metabolite graph for a small molecule structure and then generates a fragmentation graph from the metabolite graph (Figure 10). To simplify the modelling of small molecule fragmentation, we assume that mass spectrometry can only break N-C, O-C and C-C bonds. This is a reasonable assumption, as among top ten most frequent bonds in AntiMarin, these three bonds are the only single bonds that do not contain a hydrogen atom (Supplementary Table S3).

**Figure 10:**
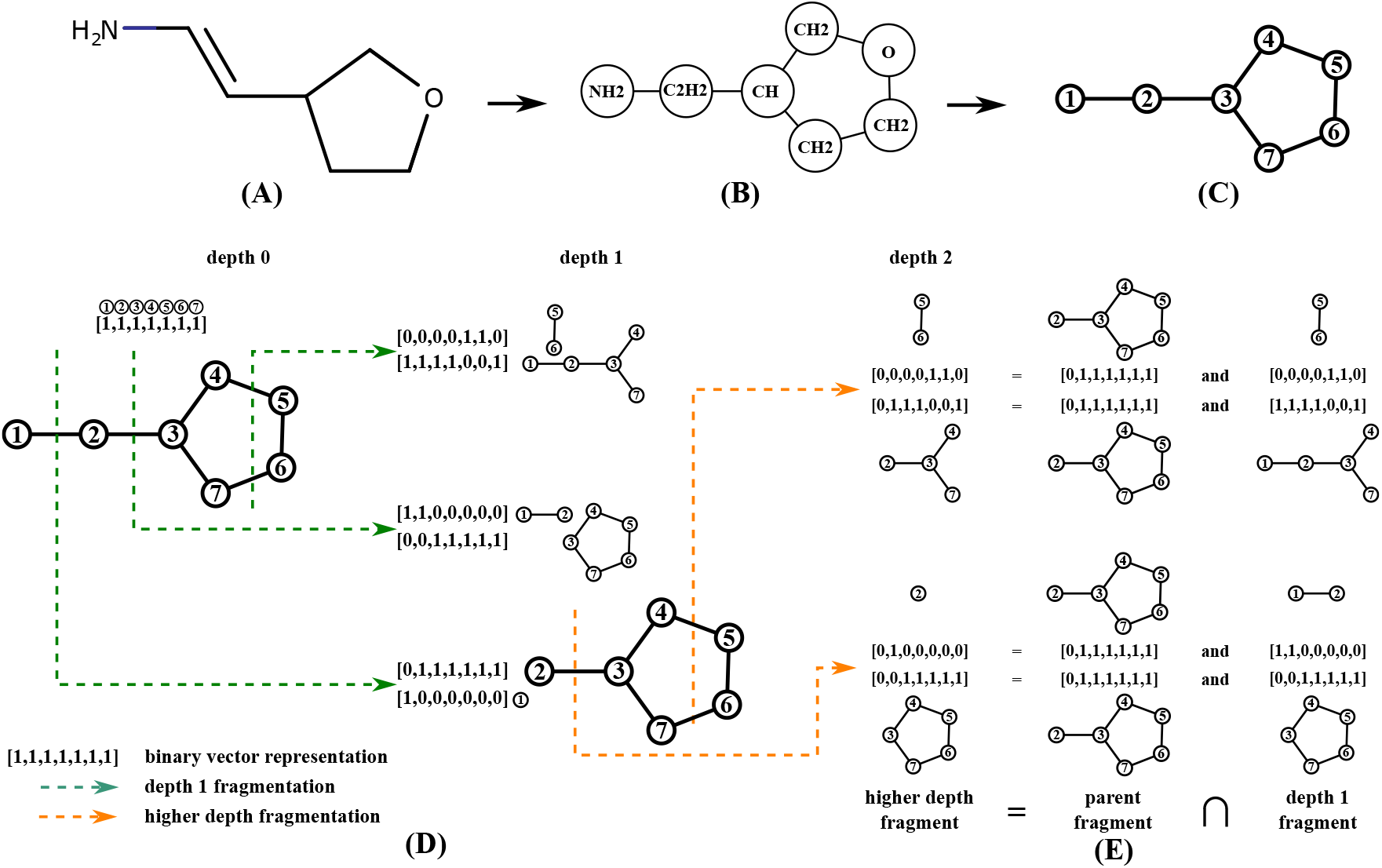
Overview of fragmentation graph construction algorithm. (A) A small molecule. (B) Metabolite graph of the molecule. (C) Indexed metabolite graph. (D) Construction of the fragmentation graph of the small molecule. We assign an index to each component of a molecule and represent each of its fragments as a binary vector, where 1 and 0 indicate presence and absence of the components. Each bridge/2-cut of a parent fragment gives two child fragments. (E) Higher depth fragments can be computed as overlaps of their parents with the depth one fragments.

To construct a metabolite graph, molDiscovery first disconnects N-C, O-C and C-C bonds. The resulting connected components form the nodes of the metabolite graph. Edges in the metabolite graph correspond to bonds between the connected components (Figure 10AB).

The fragmentation graph of a molecule is a directed acyclic graph with a single source (the metabolite graph) where nodes are fragments of the molecule and directed edges between the nodes represent bridge or 2-cut fragmentations. To construct a fragmentation graph, molDiscovery first searches for all the bridges and 2-cuts of the metabolite graph to obtain depth one fragments of the molecule using Hopcroft and Tarjan’s algorithm for graph manipulation [50]. Each fragment of the molecule can be represented by a fixed length binary vector that indicates the presence or absence of metabolite graph nodes in the fragment (Figure 10D). We observe that depth two fragments can be formed by a bitwise `AND` operation between their parent fragments and depth one fragments (Figure 10E). This generalizes to computing fragments at depth *n* > 1 by intersecting their parents (fragments at depth *n* – 1) with fragments at depth one. The final fragmentation graph is constructed by connecting the source node to all depth one fragment nodes, and then iteratively connecting depth *n* – 1 fragment nodes to the corresponding fragments at depth *n* (Supplementary Algorithm).

### Learning a probabilistic model to match mass spectra and small molecules

Dereplicator+ uses a naïve scoring scheme which ignores that (i) fragmentation probability of different bonds are different, (ii) bridges have a higher chance of fragmentation than 2-cuts, (iii) peaks with higher intensity have higher chances of corresponding to a fragmentation than lower intensity peaks, (iv) matches are biased towards molecules with large fragmentation graph size, and (v) matches are biased towards spectra with a large number of peaks.

In order to solve the above shortcomings we develop a probabilistic model for matching spectra with small molecules. Given a molecule-spectrum pair in the training data (Figure 1AB), we first construct the fragmentation graph of the molecule using our fragmentation graph construction algorithm (Figure 1C). Each fragment in the fragmentation graph is assigned a (*bondType, logRank*) label. *bondType* represents the bond(s) disconnected in the parent fragment to produce the current fragment. It can either be one bond (bridge) or two bonds (2-cut). Bridges can be OC, NC, CC, while 2-cuts can be their pairwise combinations.

*logRank* represents the intensity of the mass peak corresponding to the fragment (Figure 1CD). The mass spectrometry community has used intensity rank as an abundance measure of mass peaks in a spectrum [51, 52]. The higher the intensity rank is (closer to rank 1), the more abundant is the corresponding fragment. To reduce the number of parameters and avoid overfitting, we group peaks according to their *logRank* instead of directly using intensity rank (Supplementary Note S1). A fragment will be annotated with a *logRank* between 1 – 6 if there is a peak with rank between 1 – 64 in the spectrum within 0.01 Da of the mass of the fragment (Figure 1CD). If there is no such mass peak, the fragment will be annotated with *logRank* = 7.

In the annotated fragmentation graph (Figure 1C) we assume that (i) the *logRank* of each fragment depends only on its *bondType* and the *logRank* of its parent. Here, we only consider direct parent, as considering grandparents of the fragments increases the number of parameters by an order of magnitude, resulting in overfitting. We further assume (ii) *logRank* of each mass peak is independent from the *logRank* of other peaks. While this assumption is sometimes wrong, e.g. only one peak can have *logRank* 1, we use this assumption to simplify our probabilistic model. Finally, we assume (iii) the root node has *logRank* 0. Given a small molecule structure *Molecule* and its fragmentation graph *FG*, the probability of generating a spectrum *Spectrum* is as follows:

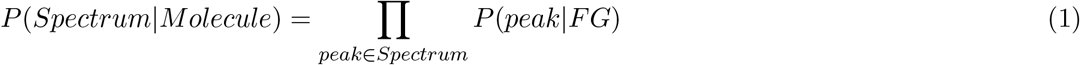

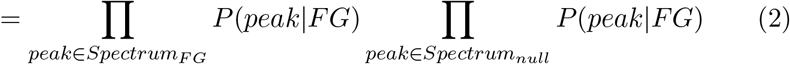

where *Spectrum_FG_* represents all the peaks within 0.01 Da of some fragment in *FG*, and *Spectrum_null_* represents the rest of the peaks. If multiple fragments match the same peak, we randomly pick one with lowest depth. Since we use *logRank* as a measure of abundance and by definition all peaks in *Spectrum_FG_* correspond to a fragment in *FG*, we can rewrite the equation as follows:

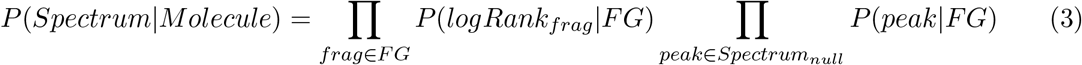

where *frag* is a fragment in the fragmentation graph. Then, by assumption (i):

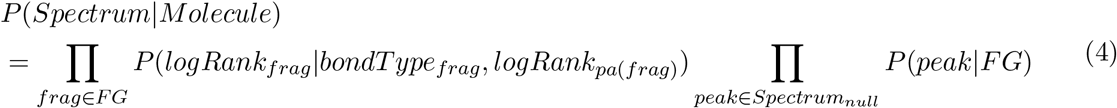

where *pa*(*frag*) is the parent of *frag*. Similarly, we can obtain the probability of generating a random spectrum (*null* model):

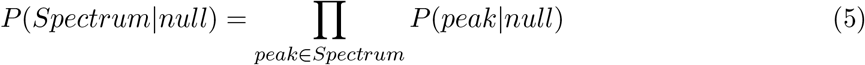

In order to learn *P*(*logRank_frag_|bondType, logRank_pa(frag)_*), we can directly count the number of the fragments with a particular *logRank* for each (*bondType* = *b,logRank_pa_* = *p*) combination in the training data as follows:

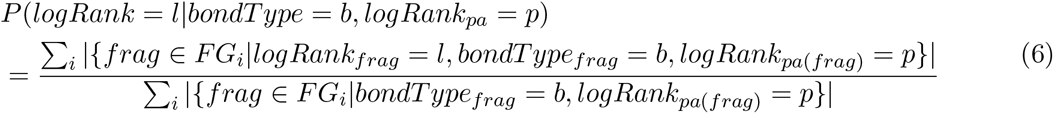

where *FG_i_* is the fragmentation graph of the *i^th^* molecule-spectrum pair in the training data.

Similarly, for the null model, we can compute the *logRank* distribution of all the mass peaks as follows:

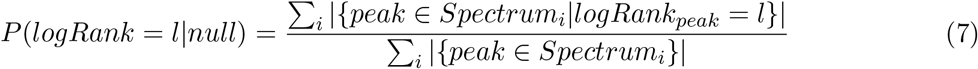

where *Spectrum_i_* is the spectrum of the *i^th^* molecule-spectrum pair.

### Scoring a spectrum against a small molecule

Given a query tandem mass spectrum (Figure 1H) and a small molecule in a chemical structure database (Figure 1G), if the spectrum precursor mass is within 0.02 Dalton of the mass of the small molecule and the user-specified adduct, they will become a candidate pair for scoring. We then construct the fragmentation graph of the molecule and annotate the query spectrum against the fragmentation graph (Figure 1IJ). Based on the described probabilistic model, we use a log-likelihood ratio model (Figure 1K) to score the spectrum against the small molecule:

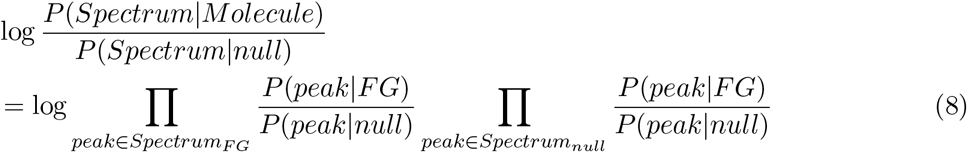

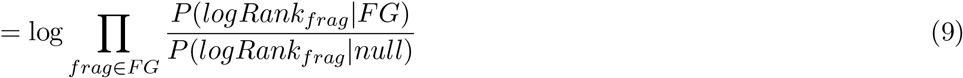

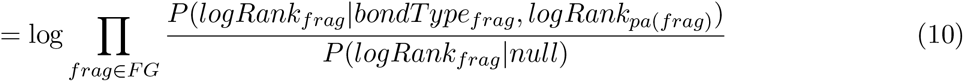

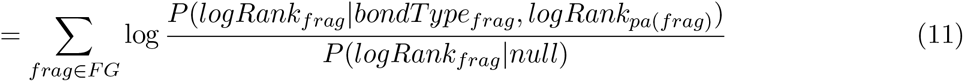

Note that in (9), we only need to consider the mass peaks corresponding to fragments in the fragmentation graph since we assume all the other peaks are generated by random chance, hence 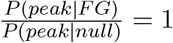.

### Computing FDR

We use target-decoy analysis to estimate FDR [53]. First, we randomly shuffle the fragmentation graph of target molecules to create decoy fragmentation graphs. Then, tandem mass spectra are searched against the target and the decoy fragmentation graph databases respectively. At each score cutoff, there are *N_target_* matches to the target database and *N_decoy_* matches to the decoy database, and the FDR is estimated as

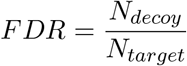

## Acknowledgement

The work of L.C., M.G., Y.L. and H.M. was supported by a research fellowship from the Alfred P. Sloan Foundation, a National Institutes of Health New Innovator Award DP2GM137413, and a U.S. Department of Energy award DE-SC0021340. The work of A.T. and A.G. was supported by RFBR (project number 20-04-01096). Research was carried out in part by computational resources provided by the Computer Center of Research park of St. Petersburg State University.

## References

[1] Rasmus Madsen, Torbjörn Lundstedt, and Johan Trygg. Chemometrics in metabolomics—a review in human disease diagnosis. Analytica chimica acta, 659(1-2):23–33, 2010.

[2] Joseph E Ippolito, Jian Xu, Sanjay Jain, Krista Moulder, Steven Mennerick, Jan R Crowley, R Reid Townsend, and Jeffrey I Gordon. An integrated functional genomics and metabolomics approach for defining poor prognosis in human neuroendocrine cancers. Proceedings of the National Academy of Sciences, 102(28):9901–9906, 2005.

[3] Ramón Estruch, Emilio Ros, Jordi Salas-Salvadó, Maria-Isabel Covas, Dolores Corella, Fernando Arós, Enrique Gómez-Gracia, Valentina Ruiz-Gutiérrez, Miquel Fiol, José Lapetra, et al. Primary prevention of cardiovascular disease with a mediterranean diet supplemented with extra-virgin olive oil or nuts. New England journal of medicine, 378(25):e34, 2018.

[4] JJ Vlaanderen, NA Janssen, G Hoek, P Keski-Rahkonen, DK Barupal, FR Cassee, I Gosens, M Strak, M Steenhof, Q Lan, et al. The impact of ambient air pollution on the human blood metabolome. Environmental research, 156:341–348, 2017.

[5] Jordi Sardans, Josep Penuelas, and Albert Rivas-Ubach. Ecological metabolomics: overview of current developments and future challenges. Chemoecology, 21(4):191–225, 2011.

[6] Susana P Gaudêncio and Florbela Pereira. Dereplication: racing to speed up the natural products discovery process. Natural product reports, 32(6):779–810, 2015.

[7] Liu Cao, Egor Shcherbin, and Hosein Mohimani. A metabolome-and metagenome-wide association network reveals microbial natural products and microbial biotransformation products from the human microbiota. Msystems, 4(4):e00387–19, 2019.

[8] Daniel McDonald, Embriette Hyde, Justine W Debelius, James T Morton, Antonio Gonzalez, Gail Ackermann, Alexander A Aksenov, Bahar Behsaz, Caitriona Brennan, Yingfeng Chen, et al. American gut: an open platform for citizen science microbiome research. Msystems, 3(3):e00031–18, 2018.

[9] Luke R Thompson, Jon G Sanders, Daniel McDonald, Amnon Amir, Joshua Ladau, Kenneth J Locey, Robert J Prill, Anupriya Tripathi, Sean M Gibbons, Gail Ackermann, et al. A communal catalogue reveals earth’s multiscale microbial diversity. Nature, 551(7681):457–463, 2017.

[10] Mingxun Wang, Jeremy J Carver, Vanessa V Phelan, Laura M Sanchez, Neha Garg, Yao Peng, Don Duy Nguyen, Jeramie Watrous, Clifford A Kapono, Tal Luzzatto-Knaan, et al. Sharing and community curation of mass spectrometry data with global natural products social molecular networking. Nature biotechnology, 34(8):828–837, 2016.

[11] Julia M Gauglitz, Christine M Aceves, Alexander A Aksenov, Gajender Aleti, Jehad Almaliti, Amina Bouslimani, Elizabeth A Brown, Anaamika Campeau, Andrés Mauricio Caraballo-Rodríguez, Rama Chaar, et al. Untargeted mass spectrometry-based metabolomics approach unveils molecular changes in raw and processed foods and beverages. Food chemistry, 302:125290, 2020.

[12] Sunghwan Kim, Jie Chen, Tiejun Cheng, Asta Gindulyte, Jia He, Siqian He, Qingliang Li, Benjamin A Shoemaker, Paul A Thiessen, Bo Yu, et al. Pubchem 2019 update: improved access to chemical data. Nucleic acids research, 47(D1):D1102–D1109, 2019.

[13] John Buckingham. Dictionary of natural products, supplement 4, volume 11. CRC press, 1997.

[14] JW Blunt, MHG Munro, and H Laatsch. Antimarin database. University of Canterbury, 432, 2006.

[15] Liu Cao, Alexey Gurevich, Kelsey L Alexander, C Benjamin Naman, Tiago Leão, Evgenia Glukhov, Tal Luzzatto-Knaan, Fernando Vargas, Robby Quinn, Amina Bouslimani, et al. Metaminer: a scalable peptidogenomics approach for discovery of ribosomal peptide natural products with blind modifications from microbial communities. Cell Systems, 9(6):600–608, 2019.

[16] Alastair W Hill and Russell J Mortishire-Smith. Automated assignment of high-resolution collisionally activated dissociation mass spectra using a systematic bond disconnection approach. Rapid Communications in Mass Spectrometry: An International Journal Devoted to the Rapid Dissemination of Up-to-the-Minute Research in Mass Spectrometry, 19(21):3111–3118, 2005.

[17] Dennis W Hill, Tzipporah M Kertesz, Dan Fontaine, Robert Friedman, and David F Grant. Mass spectral metabonomics beyond elemental formula: chemical database querying by matching experimental with computational fragmentation spectra. Analytical chemistry, 80(14):5574–5582, 2008.

[18] Sebastian Wolf, Stephan Schmidt, Matthias Müller-Hannemann, and Steffen Neumann. In silico fragmentation for computer assisted identification of metabolite mass spectra. BMC bioinformatics, 11(1):148, 2010.

[19] Ralf Gugisch, Adalbert Kerber, Axel Kohnert, Reinhard Laue, Markus Meringer, Christoph Rücker, and Alfred Wassermann. Molgen 5.0, a molecular structure generator. In Advances in mathematical chemistry and applications, pages 113–138. Elsevier, 2015.

[20] Martin Krauss, Heinz Singer, and Juliane Hollender. Lc–high resolution ms in environmental analysis: from target screening to the identification of unknowns. Analytical and bioanalytical chemistry, 397(3):943–951, 2010.

[21] Hiroshi Tsugawa, Tobias Kind, Ryo Nakabayashi, Daichi Yukihira, Wataru Tanaka, Tomáš Cajka, Kazuki Saito, Oliver Fiehn, and Masanori Arita. Hydrogen rearrangement rules: computational ms/ms fragmentation and structure elucidation using ms-finder software. Analytical chemistry, 88(16):7946–7958, 2016.

[22] Hosein Mohimani, Alexey Gurevich, Alexander Shlemov, Alla Mikheenko, Anton Korobeynikov, Liu Cao, Egor Shcherbin, Louis-Felix Nothias, Pieter C Dorrestein, and Pavel A Pevzner. Dereplication of microbial metabolites through database search of mass spectra. Nature communications, 9(1):1–12, 2018.

[23] Gert Wohlgemuth, Sajjan S Mehta, Ramón F Mejia, Steffen Neumann, Diego Pedrosa, Tomas Pluskal, Emma L Schymanski, Egon L Willighagen, Michael Wilson, David S Wishart, et al. Splash, a hashed identifier for mass spectra. Nature biotechnology, 34(11):1099–1101, 2016.

[24] Greg Landrum et al. Rdkit: Open-source cheminformatics. 2006.

[25] Jiangyong Gu, Yuanshen Gui, Lirong Chen, Gu Yuan, Hui-Zhe Lu, and Xiaojie Xu. Use of natural products as chemical library for drug discovery and network pharmacology. PloS one, 8(4):e62839, 2013.

[26] David S Wishart, Dan Tzur, Craig Knox, Roman Eisner, An Chi Guo, Nelson Young, Dean Cheng, Kevin Jewell, David Arndt, Summit Sawhney, et al. Hmdb: the human metabolome database. Nucleic acids research, 35(suppl_1):D521–D526, 2007.

[27] Manish Sud, Eoin Fahy, Dawn Cotter, Alex Brown, Edward A Dennis, Christopher K Glass, Alfred H Merrill Jr, Robert C Murphy, Christian RH Raetz, David W Russell, et al. Lmsd: Lipid maps structure database. Nucleic acids research, 35(suppl_1):D527–D532, 2007.

[28] Augustin Scalbert, Cristina Andres-Lacueva, Masanori Arita, Paul Kroon, Claudine Manach, Mireia Urpi-Sarda, and David Wishart. Databases on food phytochemicals and their health-promoting effects. Journal of agricultural and food chemistry, 59(9):4331–4348, 2011.

[29] Jeffrey A Van Santen, Grégoire Jacob, Amrit Leen Singh, Victor Aniebok, Marcy J Balunas, Derek Bunsko, Fausto Carnevale Neto, Laia Castaño-Espriu, Chen Chang, Trevor N Clark, et al. The natural products atlas: an open access knowledge base for microbial natural products discovery. ACS central science, 5(11):1824–1833, 2019.

[30] Minoru Kanehisa and Susumu Goto. Kegg: kyoto encyclopedia of genes and genomes. Nucleic acids research, 28(1):27–30, 2000.

[31] David S Wishart, Craig Knox, An Chi Guo, Savita Shrivastava, Murtaza Hassanali, Paul Stothard, Zhan Chang, and Jennifer Woolsey. Drugbank: a comprehensive resource for in silico drug discovery and exploration. Nucleic acids research, 34(suppl_1):D668–D672, 2006.

[32] Xavier Lucas, Christian Senger, Anika Erxleben, Björn A Grüning, Kersten Döring, Johannes Mosch, Stephan Flemming, and Stefan Günther. Streptomedb: a resource for natural compounds isolated from streptomyces species. Nucleic acids research, 41(D1):D1130–D1136, 2012.

[33] Marnix H Medema, Renzo Kottmann, Pelin Yilmaz, Matthew Cummings, John B Biggins, Kai Blin, Irene De Bruijn, Yit Heng Chooi, Jan Claesen, R Cameron Coates, et al. Minimum information about a biosynthetic gene cluster. Nature chemical biology, 11(9):625–631, 2015.

[34] Vanessa Neveu, Jara Perez-Jiméenez, Femke Vos, Vanessa Crespy, Lerman du Chaffaut, Louise Mennen, Craig Knox, Roman Eisner, J Cruz, D Wishart, et al. Phenol-explorer: an online comprehensive database on polyphenol contents in foods. Database, 2010, 2010.

[35] Evelien Wynendaele, Antoon Bronselaer, Joachim Nielandt, Matthias D’Hondt, Sofie Stalmans, Nathalie Bracke, Frederick Verbeke, Christophe Van De Wiele, Guy De Tré, and Bart De Spiegeleer. Quorumpeps database: chemical space, microbial origin and functionality of quorum sensing peptides. Nucleic Acids Research (submitted for publication), 2012.

[36] Kai Dührkop, Markus Fleischauer, Marcus Ludwig, Alexander A Aksenov, Alexey V Melnik, Marvin Meusel, Pieter C Dorrestein, Juho Rousu, and Sebastian Böcker. Sirius 4: a rapid tool for turning tandem mass spectra into metabolite structure information. Nature methods, 16(4):299–302, 2019.

[37] Dries Verdegem, Diether Lambrechts, Peter Carmeliet, and Bart Ghesquière. Improved metabolite identification with midas and magma through ms/ms spectral dataset-driven parameter optimization. Metabolomics, 12(6):98, 2016.

[38] Felicity Allen, Russ Greiner, and David Wishart. Competitive fragmentation modeling of esi-ms/ms spectra for putative metabolite identification. Metabolomics, 11(1):98–110, 2015.

[39] Kai Dührkop, Huibin Shen, Marvin Meusel, Juho Rousu, and Sebastian Böcker. Searching molecular structure databases with tandem mass spectra using csi: Fingerid. Proceedings of the National Academy of Sciences, 112(41):12580–12585, 2015.

[40] Yannick Djoumbou Feunang, Roman Eisner, Craig Knox, Leonid Chepelev, Janna Hastings, Gareth Owen, Eoin Fahy, Christoph Steinbeck, Shankar Subramanian, Evan Bolton, et al. Classyfire: automated chemical classification with a comprehensive, computable taxonomy. Journal of cheminformatics, 8(1):61, 2016.

[41] Don D Nguyen, Alexey V Melnik, Nobuhiro Koyama, Xiaowen Lu, Michelle Schorn, Jinshu Fang, Kristen Aguinaldo, Tommie L Lincecum, Maarten GK Ghequire, Victor J Carrion, et al. Indexing the pseudomonas specialized metabolome enabled the discovery of poaeamide b and the bananamides. Nature microbiology, 2(1):1–10, 2016.

[42] Alexey Gurevich, Alla Mikheenko, Alexander Shlemov, Anton Korobeynikov, Hosein Mohimani, and Pavel A Pevzner. Increased diversity of peptidic natural products revealed by modification-tolerant database search of mass spectra. Nature microbiology, 3(3):319–327, 2018.

[43] Stephen F Altschul, Warren Gish, Webb Miller, Eugene W Myers, and David J Lipman. Basic local alignment search tool. Journal of molecular biology, 215(3):403–410, 1990.

[44] James R Doroghazi, Jessica C Albright, Anthony W Goering, Kou-San Ju, Robert R Haines, Konstantin A Tchalukov, David P Labeda, Neil L Kelleher, and William W Metcalf. A roadmap for natural product discovery based on large-scale genomics and metabolomics. Nature chemical biology, 10(11):963, 2014.

[45] Jorge C Navarro-Muñoz, Nelly Selem-Mojica, Michael W Mullowney, Satria A Kautsar, James H Tryon, Elizabeth I Parkinson, Emmanuel LC De Los Santos, Marley Yeong, Pablo Cruz-Morales, Sahar Abubucker, et al. A computational framework to explore large-scale biosynthetic diversity. Nature chemical biology, 16(1):60–68, 2020.

[46] Li Yang, Hanxiang Li, Ping Wu, Ahmed Mahal, Jinghua Xue, Liangxiong Xu, and Xiaoyi Wei. Dinghupeptins a–d, chymotrypsin inhibitory cyclodepsipeptides produced by a soil-derived streptomyces. Journal of natural products, 81(9):1928–1936, 2018.

[47] Tilmann Weber, Kai Blin, Srikanth Duddela, Daniel Krug, Hyun Uk Kim, Robert Bruccoleri, Sang Yup Lee, Michael A Fischbach, Rolf Müller, Wolfgang Wohlleben, et al. antismash 3.0—a comprehensive resource for the genome mining of biosynthetic gene clusters. Nucleic acids research, 43(W1):W237–W243, 2015.

[48] Marc Röttig, Marnix H Medema, Kai Blin, Tilmann Weber, Christian Rausch, and Oliver Kohlbacher. Nrpspredictor2—a web server for predicting nrps adenylation domain specificity. Nucleic acids research, 39(suppl_2):W362–W367, 2011.

[49] Makoto Ubukata, Masakasu Uramoto, Jun Uzawa, and Kiyoshi Isono. Structure and biological activity of neopeptins a, b and c, inhibitors of fungal cell wall glycan synthesis. Agricultural and biological chemistry, 50(2):357–365, 1986.

[50] John Hopcroft and Robert Tarjan. Algorithm 447: efficient algorithms for graph manipulation. Communications of the ACM, 16(6):372–378, 1973.

[51] Sangtae Kim and Pavel A Pevzner. Ms-gf+ makes progress towards a universal database search tool for proteomics. Nature communications, 5:5277, 2014.

[52] Azat M Tagirdzhanov, Alexander Shlemov, and Alexey Gurevich. Nps: scoring and evaluating the statistical significance of peptidic natural product–spectrum matches. Bioinformatics, 35(14):i315–i323, 2019.

[53] Joshua E Elias and Steven P Gygi. Target-decoy search strategy for increased confidence in large-scale protein identifications by mass spectrometry. Nature methods, 4(3):207, 2007.

