## Supplemental Notes, Figures and Tables for "MolDiscovery: Learning Mass Spectrometry Fragmentation of Small Molecules"

### 1 Supplementary Note

**Supplementary Note S1. LogRank computation.** Equation (1) shows the formula used to compute the *logRank* of a peak from its *rank*. Note that multiple peaks with different ranks can be assigned to the same *logRank*. Additionally there are only 8 possible values of *logRank* ($logRank = 0$ is reserved for the root node). The number of peaks with $logRank{=}i$ is $2^{i-1}$. For example, $logRank = 1$ represents the most intense peak, $logRank = 2$ represents the next $2^1 = 2$ most intense peaks, and $logRank = 3$ represents the next $2^2 = 4$ most intense peaks. We define the lowest (farthest from rank 1) possible *logRank* as 7 and all peaks that would be assigned to a *logRank* larger then 7 are instead assigned $logRank = 7$. Note that using *logRank*, the number of parameters of the model reduces from 64 per *bondType* to only 7 per *bondType*.

$$logRank = \min(\lfloor \log_2(rank) \rfloor + 1, 7) \qquad (1)$$

Fluctuations in peak intensity due to noise could change the annotated *logRank*. However, *logRank*s are more robust to these fluctuations in compare to ranks as the slight change of peak intensity will leads to a small change (at most 1) in *logRank*. In addition, the change of probability score due to the change of *logRank* is smooth (Supplmentary Figure S1).

#### 2 Supplementary Algorithm

**Algorithm 1** Fragmentation Graph Construction Algorithm

**Input:** Metabolite graph *metGraph*, maximum depth *maxDepth*
**Output:** Fragmentation graph *FG*

$root \leftarrow [1, \ldots, 1]$ // root contains all nodes in *metGraph*
$depth_1Frags \leftarrow \{frag \mid frag \in \text{HopcroftTarjan}(metGraph)\}$
$FGNodes \leftarrow \{root\} \cup depth_1Frags$
$FGEdges \leftarrow \{(root, frag) \mid frag \in depth_1Frags\}$
$prevDepthFrags \leftarrow depth_1Frags$
**for** $i \in \{2, \ldots, maxDepth\}$ **do**
  $currDepthFrags \leftarrow \emptyset$
  **for all** $parentFrag \in prevDepthFrags$ **do**
    **for all** $depth_1Frag \in depth_1Frags$ **do**
      $newFrag \leftarrow parentFrag \,\&\, depth_1Frag$
      $currDepthFrags \leftarrow currDepthFrags \cup \{newFrag\}$
      $FGNodes \leftarrow FGNodes \cup \{newFrag\}$
      $FGEdges \leftarrow FEdges \cup \{(parentFrag, newFrag)\}$
    **end for**
  **end for**
  $prevDepthFrags \leftarrow currDepthFrags$
**end for**
**return** $(FGNodes, FGEdges)$

### 3 Supplementary Figures

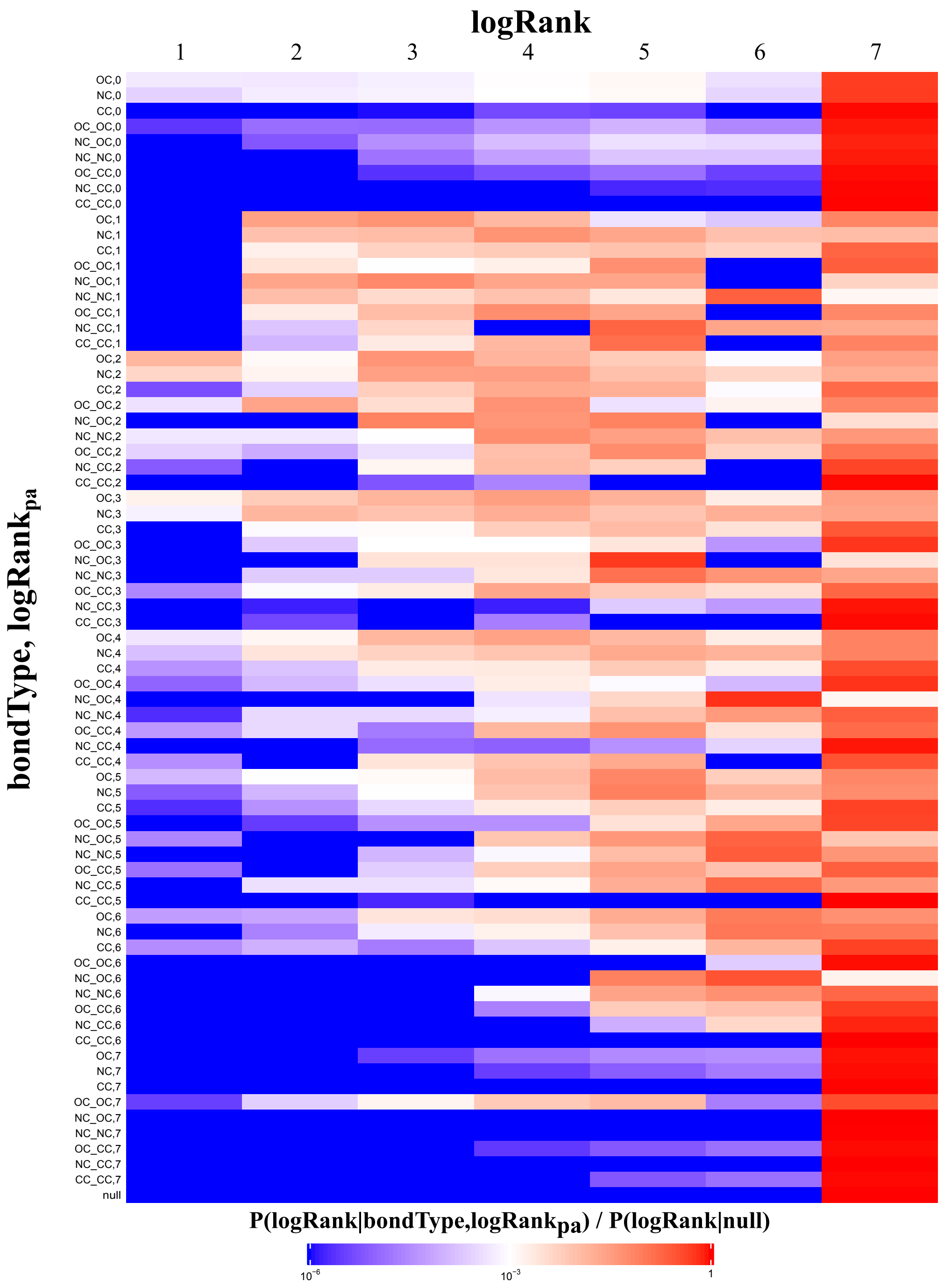


**Figure S1:** Heatmap of $P(logRank|bondType, logRank_{pa})$ for charge +1 fragments. Each row represents $bondType$ and $logRank_{pa}$. The row "null" refers to the null distribution $P(logRank|null)$. Each column represents the $logRank$ of a child fragment. When $logRank_{pa}$ is 0, it means the parent is the root (precursor molecule).

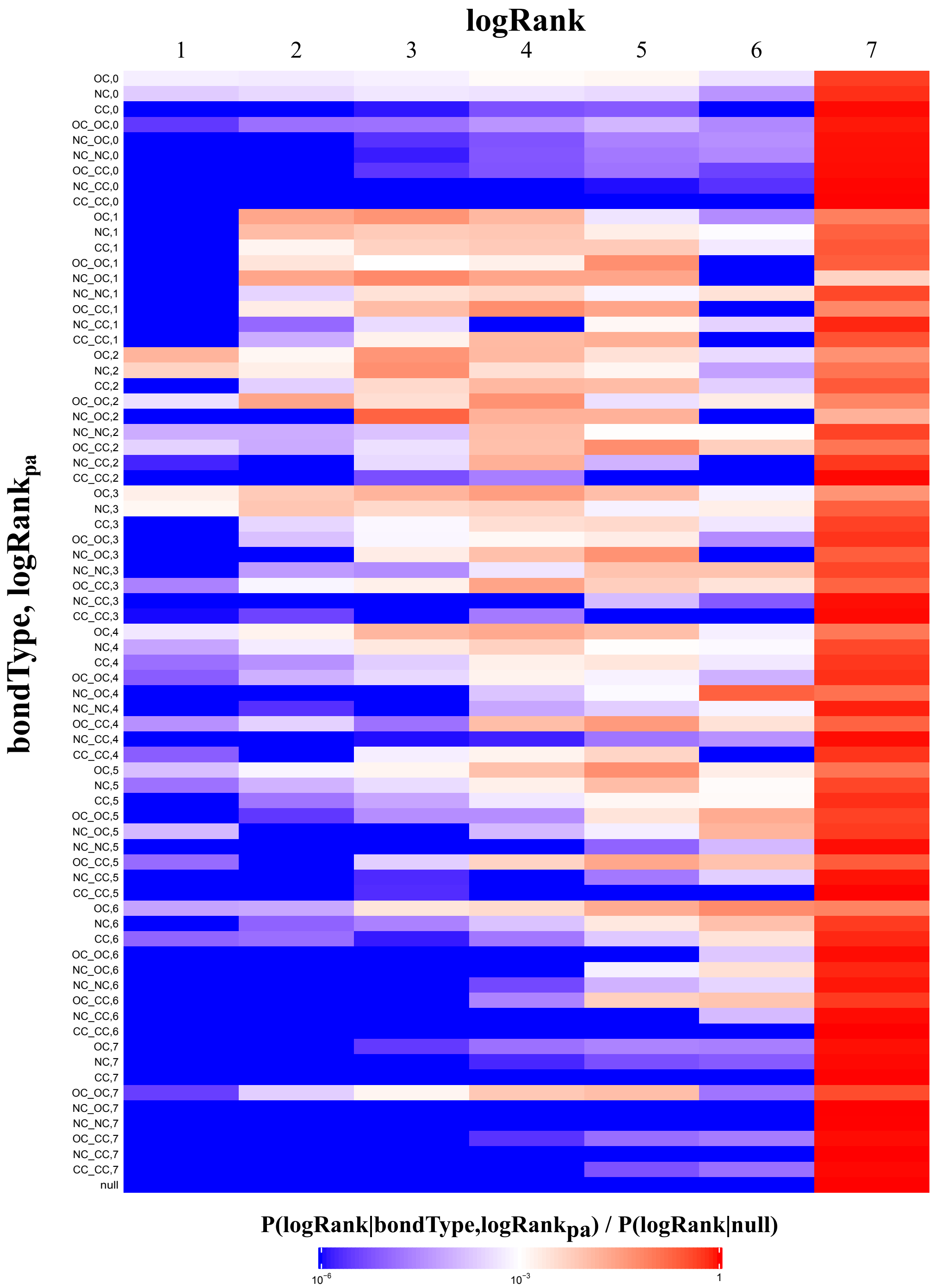


**Figure S2:** Heatmap of $P(logRank|bondType, logRank_{pa})$ for charge +2 fragments. Each row represents $bondType$ and $logRank_{pa}$. The row "null" refers to the null distribution $P(logRank|null)$. Each column represents the $logRank$ of a child fragment. When $logRank_{pa}$ is 0, it means the parent is the root (precursor molecule).

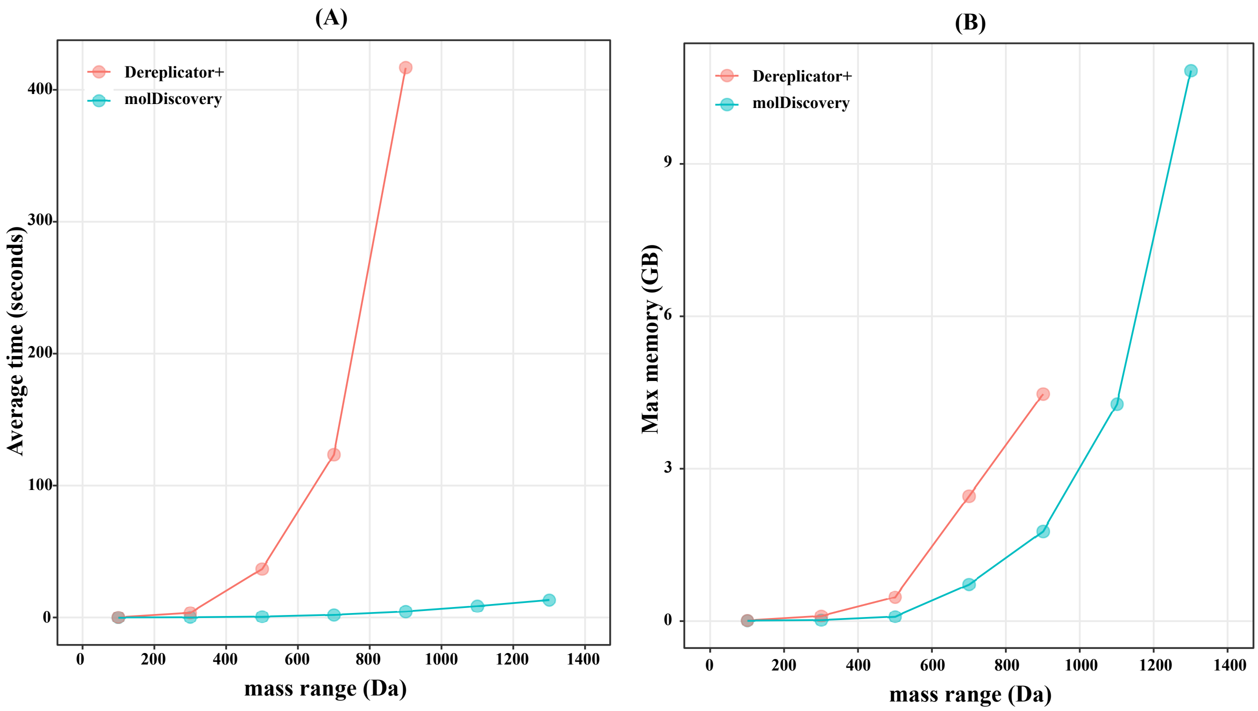


**Figure S3:** Benchmarking fragmentation tree construction algorithms. We compare the (A) average fragmentation tree construction time and (B) maximum memory (resident set size) consumption of molDiscovery with Dereplicator+ on the DNP database over molecules of various masses. Note that we randomly selected 200 molecules from the DNP database for each 200 Da mass window to compute average running time per molecule as well as maximum memory usage. For cases exceeding the memory limit of 15 GB, no data point is shown.

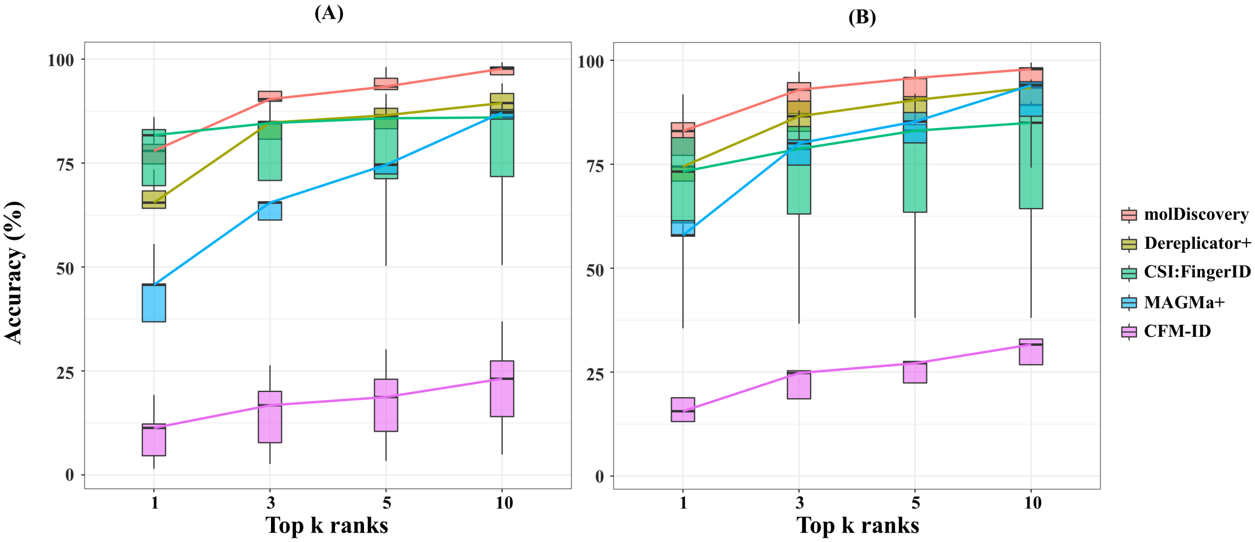


**Figure S4:** Benchmarking the database search accuracy on (A) GNPS spectral library (4,781 molecules) and (B) GNPS spectral library excluding CSI:FingerID training data (1,739 molecules). For each spectrum, we rank the matching scores given by molDiscovery, Dereplicator+, MAGMa+ 1.0.1, CFM-ID 2.0, and CSI:FingerID 1.4.3 (from SIRIUS 4.4.2). If there is a tie, we assign the average rank to all the matches in the tie. Then we check if the correct molecule-spectrum match is among the top $k$ ($k = 1, 3, 5, 10$) matches. The x-axis is the rank $k$, and the y-axis is the proportion of correct molecule-spectrum matches in the test dataset that are among the top $k$ matches. Center lines in the box plots denote the median accuracy of each method across the five cross-validation folds.

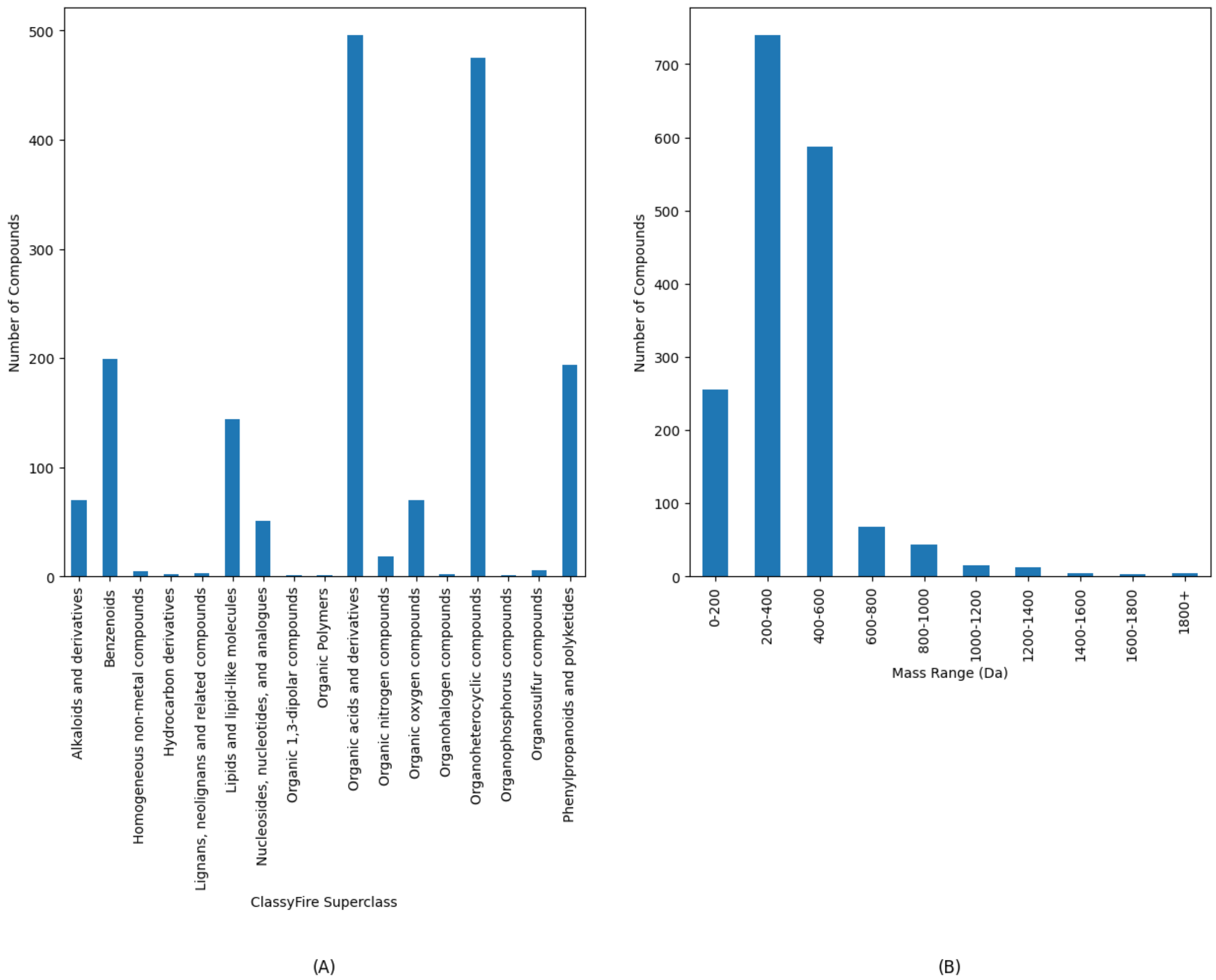


**Figure S5:** Number of compounds in each division of the non-redundant GNPS dataset. (A) Number of compounds classified as each superclass found by ClassyFire. (B) Number of compounds that fall within different mass ranges.

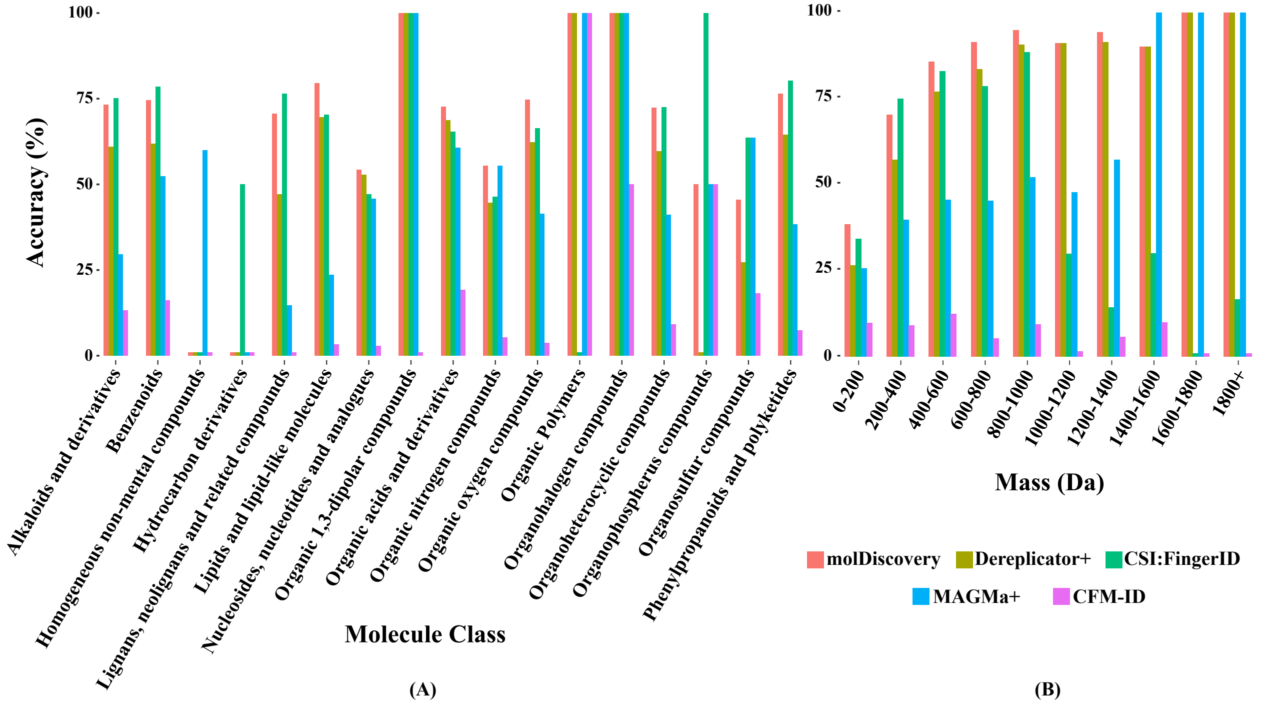


**Figure S6:** Database search accuracy of molDiscovery, Dereplicator+, CSI:FingerID, MAGMa+, and CFM-ID on molecules of (A) different ClassyFire superclasses, and (B) different mass ranges, on GNPS spectral library compounds.

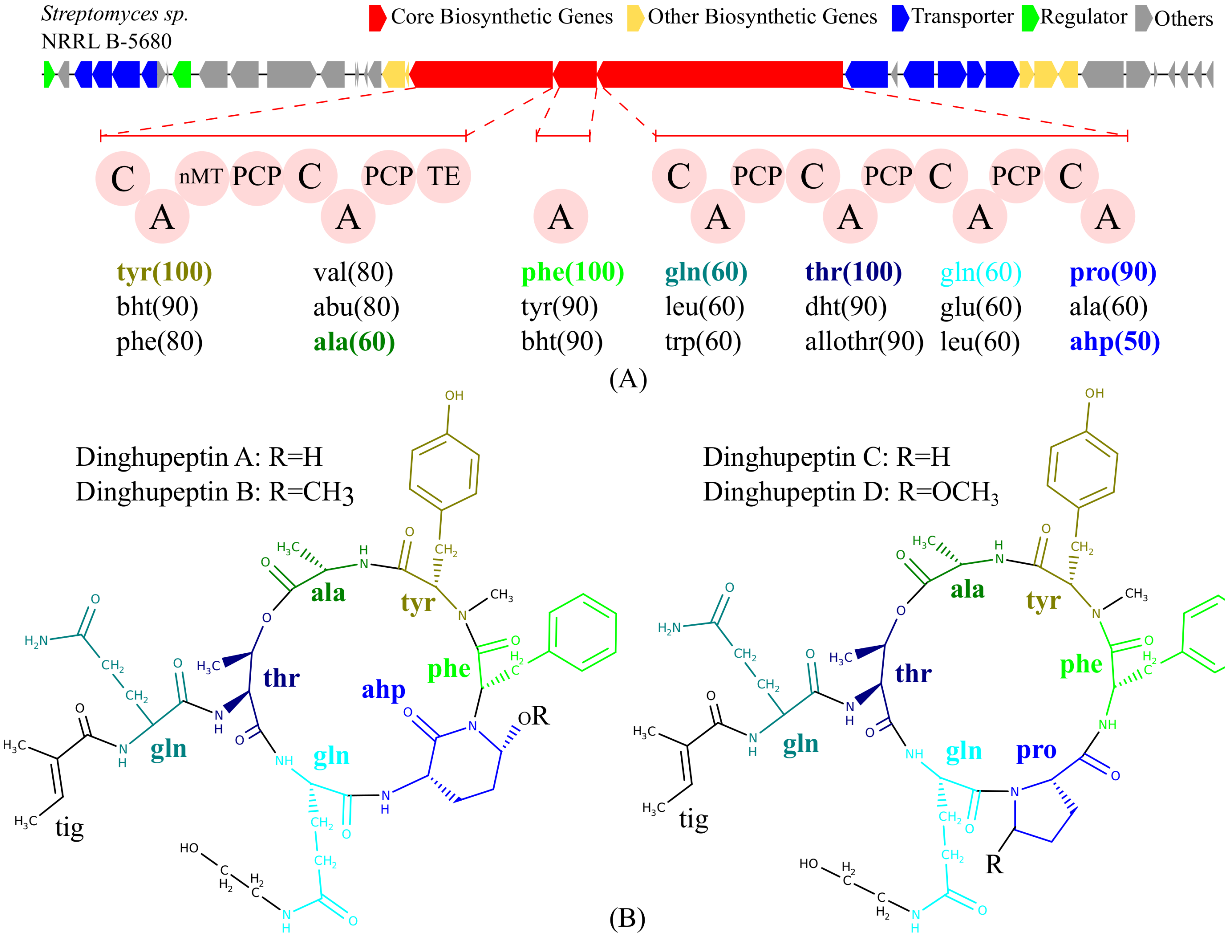


**Figure S7:** (A) Biosynthetic gene cluster of the dinghupeptin family. MolDiscovery identified dinghupeptin A at FDR 0% in *Streptomyces* NRRL B-5680. After searching its genome using antiSMASH, we detected a non-ribosomal peptide biosynthetic gene cluster with adenylation domains which show high specificity to the amino acid residues in dinghupeptin molecules. (B) Molecular structures of dinghupeptin A-D identified by molDiscovery at FDR 0.01%. Ahp stands for 3-amino-6-hydroxypiperidone and tig stands for tiglic acid.

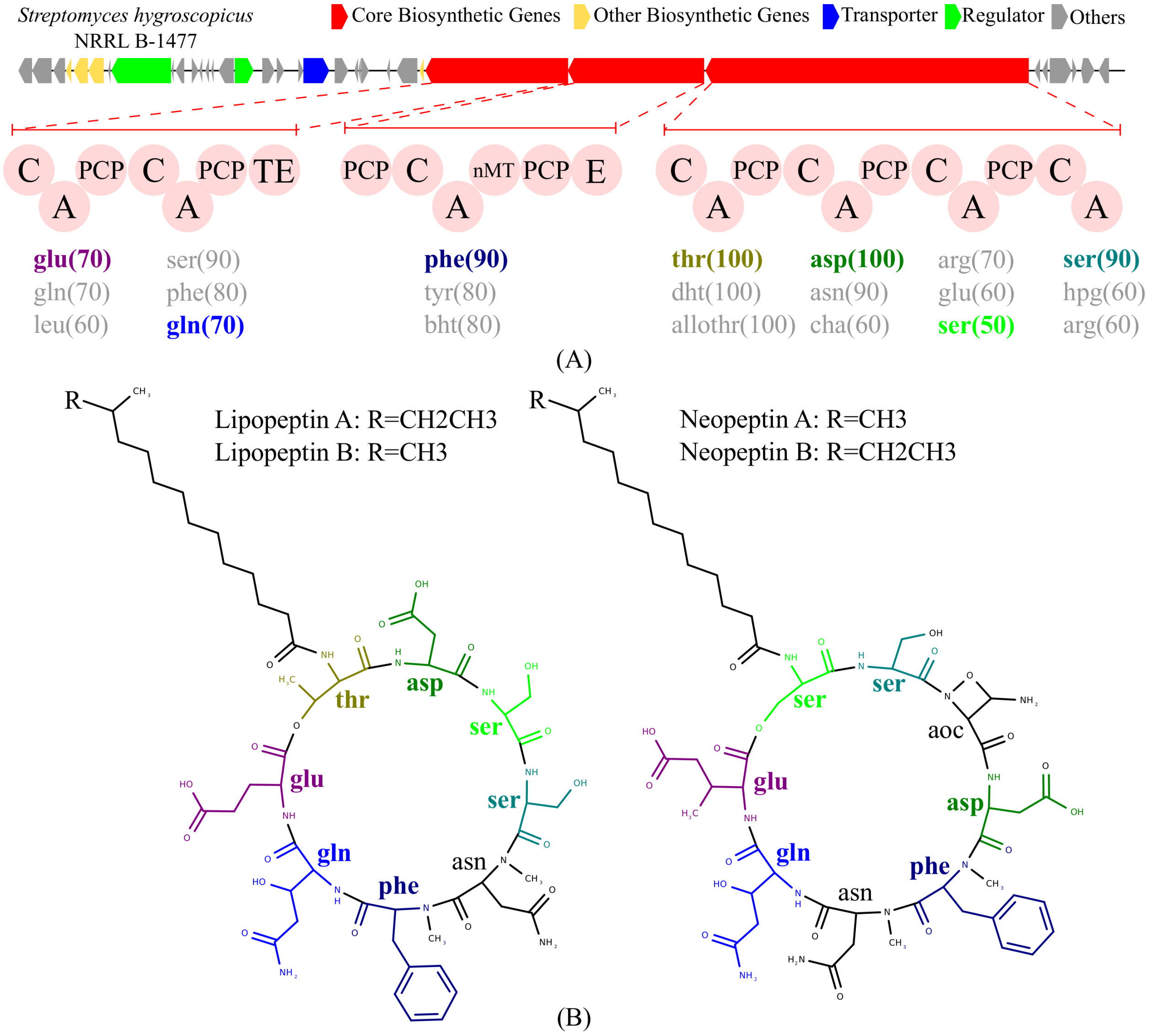


**Figure S8:** (A) Biosynthetic gene cluster of the lipopeptin and neopeptin families. MolDiscovery identified the molecules of both lipopeptin family and neopeptin family at FDR 0% in *Streptomyces hygroscopicus* NRRL B-1477. After searching its genome using antiSMASH, we detected a non-ribosomal peptide biosynthetic gene cluster with adenylation domains which show high specificity to the amino acid residues in both families. (B) Molecular Structure of lipopeptin A-B and neopeptin A-B identified by molDiscovery at FDR 0%. Aoc stands for 3-amino-2-oxazetidine-4-carboxylic acid.

### 4 Supplementary Tables.

| MassiveID | NumSpectra | MassiveID | NumSpectra |
|---|---|---|---|
| MSV000078556 | 61970 | MSV000084117 | 12287 |
| MSV000078836 | 481548 | MSV000084475 | 638641 |
| MSV000078839 | 403604 | MSV000084674 | 28854 |
| MSV000078847 | 28615 | MSV000084723 | 136043 |
| MSV000078850 | 59175 | MSV000084771 | 442 |
| MSV000078891 | 207413 | MSV000084884 | 85130 |
| MSV000078995 | 818 | MSV000084945 | 1550594 |
| MSV000079015 | 6085 | MSV000084954 | 15513 |
| MSV000079139 | 4435 | MSV000084989 | 808 |
| MSV000079284 | 27818 | MSV000085003 | 8526 |
| MSV000079519 | 49352 | MSV000085018 | 44223 |
| MSV000080251 | 1462003 | MSV000085023 | 2372 |
| MSV000080427 | 12102 | MSV000085026 | 1872 |
| MSV000081063 | 28563 | MSV000085027 | 426 |
| MSV000081318 | 6376 | MSV000085032 | 2660 |
| MSV000081504 | 607 | MSV000085123 | 769 |
| MSV000082045 | 1665897 | MSV000085158 | 25606 |
| MSV000082285 | 943 | MSV000085159 | 188737 |
| MSV000082831 | 36566 | MSV000085179 | 7853 |
| MSV000083081 | 1389 | MSV000085180 | 9095 |
| MSV000083295 | 3850 | MSV000085192 | 1634 |
| MSV000083648 | 7 | MSV000085214 | 3284 |
| MSV000083734 | 289693 | MSV000083738 | 409245 |

**Table S1:** 46 GNPS spectral datasets analyzed in the paper. In each dataset we only analyze the mass spectral files that have corresponding genome assemblies. There are 8,013,433 spectra in total.

| Molecule | MSV ID | Score | MZ | RT | Adduct |
|---|---|---|---|---|---|
| massetolide_J | MSV000085018 | 245.834 | 1112.68 | 1536.35 | M+H |
| massetolide_H | MSV000085018 | 236.383 | 1154.73 | 1745.36 | M+H |
| massetolide_I | MSV000085018 | 232.237 | 1112.68 | 1506.32 | M+H |
| Massetolide_D | MSV000085018 | 226.163 | 1140.71 | 1675.35 | M+H |
| viscosin | MSV000085018 | 221.532 | 1126.69 | 1505.12 | M+H |
| Massetolide_E | MSV000085018 | 221.267 | 1112.68 | 1559.68 | M+H |
| Massetolide_F | MSV000085018 | 221.168 | 1126.69 | 1615.11 | M+H |
| Ornibactin_C4 * | MSV000084945 | 212.54 | 737.404 | 176.698 | M+H |
| Taxlllaid_A | MSV000081063 | 209.695 | 808.549 | 627.514 | M+H |

*Ornibactin_C4_N5-Deacyl,_N5-(3-hydroxyoctanoyl)

| | | | | | |
|---|---|---|---|---|---|
| Surugamide_G | MSV000083738 | 206.771 | 884.594 | 2615.94 | M+H |
| Esperin | MSV000084117 | 206.03 | 1036.69 | 342.024 | M+H |
| Actinomycin_monolactone | MSV000083738 | 205.893 | 1273.65 | 3499.08 | M+H |
| Surugamide_C | MSV000079519 | 204.486 | 898.61 | 285.818 | M+H |
| Ornibactin_C8 | MSV000084945 | 203.704 | 737.404 | 177.16 | M+H |
| Surugamide_B | MSV000083738 | 203.433 | 898.611 | 2648.58 | M+H |
| Champacyclin | MSV000079519 | 202.923 | 898.611 | 286.793 | M+H |
| Taxlllaid_F | MSV000081063 | 202.368 | 826.56 | 545.308 | M+H |
| Gacamide_A | MSV000085018 | 197.487 | 1394.84 | 1528.63 | M+H |
| Surugamide_D | MSV000083738 | 196.059 | 898.61 | 2628.42 | M+H |
| Taxlllaid_C | MSV000081063 | 195.483 | 794.533 | 612.231 | M+H |
| Sameuramide | MSV000085192 | 192.685 | 1016.55 | 255.397 | M+H |
| Antrimycin_Antrimycin_D | MSV000083738 | 191.778 | 728.394 | 1949.09 | M+H |
| Antibiotic_MA_026_Antibiotic_MA_026 | MSV000085018 | 189.546 | 888.547 | 1789.11 | M+2H |
| Taxillaid_structure_0 | MSV000081063 | 188.787 | 808.549 | 631.648 | M+H |
| MOL1 † | MSV000085192 | 187.785 | 1002.54 | 260.242 | M+H |
| callipeltin_B | MSV000083738 | 186.724 | 1031.54 | 3525.33 | M+H |
| TG(16:0/20:1(11Z)/18:2(9Z,12Z)) | MSV000083738 | 184.924 | 883.776 | 3829.81 | M+H |
| a-Substance_Ib | MSV000084945 | 184.139 | 686.399 | 247.877 | M+H |
| surugamides_A | MSV000084723 | 183.976 | 912.621 | 579.044 | M+H |
| Anabaenopeptin_NZ857 | MSV000078891 | 183.617 | 858.435 | 624.713 | M+H |
| Ambactin | MSV000081063 | 183.167 | 751.409 | 395.928 | M+H |
| gageostatin_B | MSV000084117 | 182.802 | 1054.7 | 330.423 | M+H |
| FR_900359_FR_900359 | MSV000085192 | 182.698 | 1002.55 | 249.025 | M+H |
| xantholysin_A_structure_0 | MSV000085018 | 182.298 | 888.547 | 1476.12 | M+2H |
| Xenobovid_B | MSV000081063 | 182.022 | 794.535 | 608.085 | M+H |
| Taxlllaid_E | MSV000081063 | 181.833 | 780.519 | 583.122 | M+H |
| surugamides_C | MSV000084723 | 181.798 | 898.609 | 565.331 | M+H |
| Anabaenopeptin_NZ825_4'-Hydroxy | MSV000078891 | 180.702 | 842.44 | 682.585 | M+H |
| Xentrivalpeptides_Xentrivalpeptide_A | MSV000081063 | 180.643 | 860.487 | 558.542 | M+H |
| Xentrivalpeptides_Xentrivalpeptide_N | MSV000081063 | 180.392 | 874.502 | 580.245 | M+H |
| Xentrivalpeptide_N | MSV000081063 | 180.015 | 874.501 | 577.4 | M+H |
| Champacyclin | MSV000084723 | 179.251 | 912.621 | 579.572 | M+H |
| Anabaenopeptin_NZ841 | MSV000078891 | 178.609 | 842.44 | 687.292 | M+H |
| Insulapeptolide_F_Insulapeptolide_F | MSV000084884 | 178.424 | 1007.52 | 237.998 | M+H |
| Actinomycin_D ‡ | MSV000083738 | 175.565 | 1269.62 | 3784.46 | M+H |
| Surugamide_A | MSV000084723 | 175.405 | 912.622 | 583.592 | M+H |
| BK_10_BK_101B | MSV000080251 | 175.235 | 1035.71 | 549.998 | M+H |
| Surugamide_A | MSV000084723 | 174.85 | 912.621 | 584.585 | M+H |
| Taxlllaid_B | MSV000081063 | 173.898 | 858.528 | 570.302 | M+H |

†3-acetamido-22-benzyl-10-¡1¡(3-hydroxy-4-methyl-2-propionamidopentanoyl)oxy¿-2-methylpropyl¿-4-isopropyl-7-(1-methoxyethyl)-19-methylene-8,13,14,16,20-pentamethyl-1,5-dioxa-8,11,14,17,20-pentaazacyclodocosane-2,6,9,12,15,18,21-heptone—3-acetamido-22-benzyl-10-1[(3-hydroxy-4-methyl-2-propionamidopentanoyl)oxy]-2-methylpropyl-4-isopropyl-7-(1-methoxyethyl)-19-methylene-8,13,14,16,20-pentamethyl-1,5-dioxa-8,11,14,17,20-pentaazacyclodocosane-2,6,9,12,15,18,21-heptone.

‡Actinomycin_D,_8CI,_9CI,_JAN_43beta-Oxo

| | | | | | |
|---|---|---|---|---|---|
| Surugamide_C | MSV000084723 | 173.289 | 898.605 | 569.502 | M+H |
| Monamycin-B3 | MSV000083738 | 173.248 | 678.417 | 3688.46 | M+H |
| Lonicatenamycin | MSV000083738 | 173.163 | 777.371 | 2858.14 | M+H |
| Szentiamide_Szentiamide | MSV000081063 | 171.439 | 838.41 | 509.547 | M+H |
| Taxlllaid_D | MSV000081063 | 171.335 | 828.519 | 610.576 | M+H |
| Surugamide_H | MSV000083738 | 170.709 | 870.579 | 2548.75 | M+H |
| Oscillapeptilide_97-B | MSV000084884 | 170.623 | 1032.54 | 253.247 | M+H |
| Ogipeptin_A | MSV000079519 | 168.914 | 955.597 | 269.902 | M+H |
| Xentrivalpeptide_O | MSV000081063 | 168.301 | 846.473 | 545.671 | M+H |
| Alterochromide_A | MSV000084884 | 167.987 | 752.355 | 203.68 | M+H |
| haprolid | MSV000085018 | 167.732 | 683.441 | 2503.76 | M+H |
| TG(16:0/20:3n6/18:2(9Z,12Z)) | MSV000083738 | 167.657 | 879.743 | 3931.22 | M+H |
| Neopeptin_Neopeptin_B | MSV000083738 | 167.366 | 1190.59 | 3346.9 | M+H |
| Alterochromide_A | MSV000084475 | 166.73 | 766.373 | 138.103 | M+H |
| Nicrophorusamide_A | MSV000083738 | 166.111 | 790.402 | 2414.23 | M+H |
| Xentrivalpeptide_A | MSV000081063 | 165.515 | 430.749 | 560.037 | M+2H |
| Orfamide_A | MSV000085018 | 164.664 | 648.424 | 1876.81 | M+2H |
| FR_900359 | MSV000085192 | 164.395 | 1002.55 | 256.93 | M+H |
| Gageostatin_B | MSV000084117 | 164.099 | 1054.7 | 330.516 | M+H |
| xantholysin_B_structure_1 | MSV000085018 | 163.966 | 881.538 | 1412.8 | M+2H |
| Bacirtsin-3 § | MSV000084117 | 163.755 | 1022.68 | 338.315 | M+H |
| TG(18:1(9Z)/18:0/18:2(9Z,12Z)) | MSV000083738 | 163.401 | 883.776 | 3789.02 | M+H |
| Xentrivalpeptides_Xentrivalpeptide_L | MSV000081063 | 162.399 | 826.502 | 551.856 | M+H |
| Orfamide_B | MSV000085018 | 161.879 | 641.417 | 1809.2 | M+2H |
| YM_254890_3”-O-Deacyl | MSV000083738 | 161.501 | 789.405 | 2119.38 | M+H |
| Tuberactinomycin_O | MSV000083738 | 161.208 | 670.339 | 197.318 | M+H |
| Xenobovid_A | MSV000081063 | 160.975 | 766.503 | 562.947 | M+H |
| Xenobovid_C | MSV000081063 | 160.835 | 822.564 | 662.551 | M+H |
| Actinomycin_F9_Actinomycin_F9 | MSV000083738 | 160.523 | 1229.62 | 3641.07 | M+H |
| Monamycin-F | MSV000083738 | 159.132 | 706.449 | 3742.82 | M+H |
| Noursamycin_D | MSV000083738 | 158.697 | 776.388 | 2377.33 | M+H |
| Nostopeptolide_A1 | MSV000078891 | 158.509 | 1081.59 | 686.661 | M+H |
| largamide_A_methyl_ester | MSV000080251 | 158.361 | 856.443 | 200.125 | M+H |
| Rhabdopeptide_5 | MSV000081063 | 158.182 | 800.593 | 537.293 | M+H |
| Cherimolacyclopeptide_D | MSV000084117 | 157.621 | 653.359 | 165.687 | M+H |
| Lyngbyastatin_5 | MSV000083738 | 157.342 | 1057.49 | 2165.29 | M+H |
| Alterochromide_A’ | MSV000084475 | 157.117 | 766.373 | 137.261 | M+H |
| Mikamycin_B | MSV000085018 | 155.536 | 434.205 | 992.012 | M+2H |
| Prexenocoumacin ¶ | MSV000081063 | 155.494 | 726.379 | 435.942 | M+H |
| Paromomycin,_BAN,_INN_1-N-Ac | MSV000083738 | 155.467 | 658.314 | 119.287 | M+H |
| Actinomycin_X0$\delta$ | MSV000083738 | 154.95 | 1271.63 | 3639.15 | M+H |
| Lipopeptin-B | MSV000083738 | 154.558 | 1163.59 | 3369.37 | M+H |
| Antibiotic_LL-BM_547alpha | MSV000083738 | 154.272 | 558.239 | 178.786 | M+H |

§ Bacirtsin-3;_Bacircine_4;_Surfactin_B2

¶ Prexenocoumacin_A_N2”-Deacyl,_N2”-(4-phenylbutanoyl)

| | | | | | |
|---|---|---|---|---|---|
| BK_10_!BK_101C | MSV000080251 | 154.187 | 1035.71 | 548.596 | M+H |
| WS-9320-A | MSV000083738 | 153.781 | 1037.49 | 2342.37 | M+H |
| cyclotheonamide_E4 | MSV000083738 | 152.951 | 839.47 | 2498.38 | M+H |
| Octaminomycin_A | MSV000083738 | 152.487 | 1001.57 | 3501.35 | M+H |
| APD_I_component_a;_Surfactin_A1 | MSV000084117 | 152.247 | 1008.66 | 335.868 | M+H |
| MOL2 [‖] | MSV000080251 | 151.863 | 652.405 | 200.541 | M+H |
| Aurantin_II | MSV000083738 | 151.526 | 1255.64 | 3766.95 | M+H |
| actinomycin_X0$\delta$ | MSV000083738 | 151.301 | 1271.63 | 3522.44 | M+H |
| Pyoverdin-QH-Ld'AtAo'*-GLU | MSV000084884 | 150.477 | 1021.46 | 214.964 | M+H |
| Tuberactinamine_N | MSV000083738 | 150.162 | 542.244 | 178.021 | M+H |
| Micropeptin_SD999_Micropeptin_SD999 | MSV000084475 | 149.953 | 1000.55 | 228.416 | M+H |
| Noursamycin_E | MSV000083738 | 149.598 | 810.37 | 2407.52 | M+H |
| Azetomycin [**] | MSV000083738 | 149.51 | 1241.62 | 3538.83 | M+H |
| Alterochromide_B | MSV000083738 | 148.841 | 778.374 | 1881.97 | M+H |
| Actinomycin_D [††] | MSV000083738 | 148.296 | 1271.63 | 3542.33 | M+H |
| Torularhodin_16'-Alcohol | MSV000084954 | 148.223 | 551.423 | 312.446 | M+H |
| Heterobactin_A | MSV000083738 | 148.005 | 599.211 | 2154.92 | M+H |
| Xentrivalpeptide_Q | MSV000081063 | 147.854 | 761.42 | 524.966 | M+H |
| Nicrophorusamide_B | MSV000083738 | 146.698 | 774.41 | 2418.83 | M+H |
| Alloxanthin_Alloxanthin | MSV000084954 | 146.611 | 565.401 | 283.176 | M+H |
| PFPGPI;Halo-toxin | MSV000080251 | 146.306 | 627.35 | 232.316 | M+H |
| Dactinomycin | MSV000083738 | 146.058 | 1255.64 | 3969.03 | M+H |
| citrusin_IX | MSV000084884 | 145.893 | 754.407 | 309.98 | M+H |
| Cyanopeptolin_CP978 | MSV000083738 | 145.823 | 978.489 | 2267.88 | M+H |
| (E)-3',4'-didehydro-$\beta$,$\psi$-caroten-16'-ol | MSV000084954 | 145.624 | 551.423 | 311.062 | M+H |
| Circulocin_gamma | MSV000083738 | 145.46 | 939.627 | 2619.83 | M+H |
| actinomycin_D | MSV000083738 | 145.227 | 1255.64 | 3831.26 | M+H |
| Octaminomycin_B | MSV000083738 | 145.141 | 987.555 | 3492.43 | M+H |
| Lipodepsipeptides_KMM_1364D | MSV000084117 | 144.897 | 1050.71 | 347.18 | M+H |
| Actinomycin_D [‡‡] | MSV000083738 | 144.45 | 1227.6 | 3534.66 | M+H |
| 3$\alpha$-(4-oxo-l-proline)-actinomycin_D | MSV000084723 | 143.748 | 1269.61 | 645.196 | M+H |
| Pullularin_C | MSV000083738 | 143.71 | 762.399 | 2434.54 | M+H |
| Konbamid | MSV000081318 | 143.071 | 877.38 | 699.772 | M+H |
| Micropeptin_88B_Micropeptin_88B | MSV000083738 | 142.835 | 1079.52 | 2158.69 | M+H |
| Crocoxanthin | MSV000084954 | 142.812 | 551.423 | 310.387 | M+H |
| Ile7-Surfactin_C13 | MSV000084117 | 142.659 | 1008.66 | 335.605 | M+H |
| Orfamide_A_Orfamide_A | MSV000085018 | 142.224 | 1295.84 | 1770.94 | M+H |
| Orfamide_C_structure_2 | MSV000085018 | 142.121 | 1267.81 | 1737.82 | M+H |
| Monamycins_Monamycin_C | MSV000083738 | 142.009 | 692.433 | 3700.64 | M+H |
| Rhabdopeptide_6 | MSV000081063 | 141.754 | 814.611 | 550.31 | M+H |
| LL-BM-547a;_Tuberactinamine_A | MSV000083738 | 141.455 | 558.24 | 286.037 | M+H |

[‖] L-Valyl-L-leucyl-L-prolyl-L-valyl-L-prol

[**] Azetomycin_I;_3A(3B)-(2-Acetidinecarboxy

[††] Actinomycin_D,_8CI,_9CI,_JAN_43betaR-Hydroxy

[‡‡] Actinomycin_D,_8CI,_9CI,_JAN_3alpha,3beta-Bis(2-azetidinecarboxylic_acid)_homologue

| | | | | | |
|---|---|---|---|---|---|
| Aurantin_III | MSV000083738 | 141.389 | 1241.62 | 3600 | M+H |
| Monadoxanthin_3'-Deoxy | MSV000084954 | 141.02 | 551.423 | 311.472 | M+H |
| Pristinamycin_IC;_Vernamycin_Bg | MSV000085018 | 140.917 | 853.387 | 912.905 | M+H |
| Diperamycin | MSV000083738 | 140.658 | 857.46 | 3574.34 | M+H |
| Nostopeptolide_A3 | MSV000078891 | 140.412 | 1081.59 | 695.337 | M+H |
| YM-47141 | MSV000083738 | 140.28 | 935.447 | 2790.24 | M+H |

**Table S2:** Top 100 molDiscovery identifications in the 46 datasets (Table S1).

| bond type | all | marine | terrestial |
|---|---|---|---|
| C-H | 2050594(45.4%) | 196820(45.6%) | 1853774(45.4%) |
| C-C | 1283757(28.4%) | 121283(28.1%) | 1162474(28.4%) |
| C-O | 356218(7.8%) | 34168(7.9%) | 322050(7.8%) |
| C=C | 222100(4.9%) | 23698(5.4%) | 198402(4.9%) |
| C-N | 183013(4.0%) | 16505(3.8%) | 166508(4.0%) |
| C=O | 140625(3.1%) | 11868(2.7%) | 128757(3.1%) |
| O-H | 138568(3.0%) | 14552(3.3%) | 124016(3.0%) |
| N-H | 82728(1.8%) | 6130(1.4%) | 76598(1.8%) |
| C=N | 11747(0.2%) | 1109(0.2%) | 10638(0.2%) |
| C-S | 9179(0.2%) | 998(0.2%) | 8181(0.2%) |
| C-Br | 7998(0.1%) | 1403(0.3%) | 6595(0.1%) |
| C-Cl | 6191(0.1%) | 824(0.1%) | 5367(0.1%) |
| S=O | 5007(0.1%) | 447(0.1%) | 4560(0.1%) |

**Table S3:** Bond type frequencies in AntiMarin database. All, marine and terrestial stands for all, marine and terrestial compounds in AntiMarin database respectively. Among top 9 most frequent bonds in natural products, only C-C, C-O and C-N do not have hydrogen or double bond. Moreover, as the frequencies of C-S, C-Br and C-Cl are less than 0.2%, and there is few training data in GNPS spectral library for these bond types, we only focus on C-C, C-O and C-N in molDiscovery. These statistics is from Supplementary Table 4 of Dereplicator+.
